# Machine learning assisted health viability assay for mouse embryos with artificial confocal microscopy (ACM)

**DOI:** 10.1101/2023.07.30.550591

**Authors:** Neha Goswami, Nicola Winston, Wonho Choi, Nastasia Z. E. Lai, Rachel B. Arcanjo, Xi Chen, Nahil Sobh, Romana A. Nowak, Mark A. Anastasio, Gabriel Popescu

## Abstract

The combination of a good quality embryo and proper maternal health factors promise higher chances of a successful in vitro fertilization (IVF) procedure leading to clinical pregnancy and live birth. Of these two factors, selection of a good embryo is a controllable aspect. The current gold standard in clinical practice is visual assessment of an embryo based on its morphological appearance by trained embryologists. More recently, machine learning has been incorporated into embryo selection “packages”. Here, we report a machine-learning assisted embryo health assessment tool utilizing a quantitative phase imaging technique called artificial confocal microscopy (ACM). We present a label-free nucleus detection method with novel quantitative embryo health biomarkers. Two viability assessment models are presented for grading embryos into two classes: healthy/intermediate (H/I) or sick (S) class. The models achieve a weighted F1 score of 1.0 and 0.99 respectively on the in-distribution test set of 72 fixed embryos and a weighted F1 score of 0.9 and 0.95 respectively on the out-of-distribution test dataset of 19 time-instances from 8 live embryos.

## 1. Introduction

Predictive and efficient viability assessment is essential to identify embryos with the highest potential for implantation and ongoing development^1, 2^. Conventional methods of embryo grading involve human decision-making, where trained embryologists assign a grade to the embryo based on optical microscopy images^3^. However, this approach carries the risk of subjectivity since the information conveyed by such qualitative images is minimal, limited to only structural information and human bias is unavoidable.

Recent technological advancements in microscopy, image processing methods, and machine learning have paved the way for a new family of embryo-viability assessment tools^4, 5^. One example is the development of time-lapse incubator systems (TLS) ^6, 7^, where a camera is placed inside the incubation system and can continually monitor embryo development in real time. Time-lapse systems have been reported to provide morphokinetic markers^8^ for embryo assessments, some of which purportedly correlate with molecular marker studies^9^. However, as with traditional microscopy, the morphological information obtained, and subsequent viability assessment made is limited by the type of imaging system employed in the incubator.

While most embryo grading tools are based on the morphology of the embryo, a few studies also involve the assessment of specific proteins^10, 11^ and genetic factors in various regions of the embryo. These analyses require specialized equipment and personnel as well as methodology standardization. Live birth prediction in mouse embryos using nuclear shape/size descriptors obtained through fluorescence microscopy was reported to achieve a classification accuracy of 83.87 %^12^. However, this approach is not feasible in the clinical setting due to the use of embryo staining, which combined with fluorescence microscopy for nucleus identification carry high risks of damage to the embryo.

The studies to date strongly indicate that a combination of structural (size of the embryo and number of cells) and compositional (protein/DNA concentration) information provides a better predictor of embryo viability and subsequent implantation potential^10^. Combining both types of measurement in one, non-invasive, label-free technique could therefore help to improve the performance of machine learning algorithms for embryo viability assessment.

Machine learning is rapidly evolving in the field of embryology^13–18^ and has the potential to provide a combined assessment system. Previous studies have predominantly relied on the use of standard light microscopy to obtain images for analysis, but optically thick samples such as embryos and organoids induce higher order scattering of incident light. This makes them difficult to image with conventional light microscopy and thus limits the amount of information that qualitative optical images can convey. Specialized microscopy techniques such as non-linear and multiphoton microscopy have been the methods of choice to achieve better penetration depth and depth sectioning in such highly scattering samples^19^. However, non-linear microscopies, due to their inherent principle of operation, always involve higher excitation power, which poses a risk of photodamage^20^. Moreover, with the exception of autofluorescence-based techniques^21^ and non-linear microscopies based on harmonic generation^22^, most of these microscopies are qualitative and require external stains to be added to the sample. The addition of any external reagent not only carries a risk of chemical toxicity but may also cause perturbation of the inherent natural state of the sample microenvironment, which in turn can potentially alter the measurements obtained. Quantitative phase imaging (QPI) ^23, 24^ is a solution to the problems of photodamage and stain-perturbations.

QPI is a label-free imaging method in which the optical phase delay of incident light from the sample is extracted without adding any external reagents^24^. The optical phase delay is a significant intrinsic marker of any sample as it provides information about the refractive index fluctuations and the structural distributions of the sample. QPI has been successfully applied to various realms of biomedical research^23, 25–29^.

Gradient light interference microscopy (GLIM) is one such QPI technique that can enable imaging of highly scattering samples like embryos and spheroids^30^. GLIM is based on the principles of phase-shifting interferometry^31^ and is developed as an add-on to a standard differential interference contrast (DIC) microscope. Recently, laser-scanning confocal microscopy (LSM) was combined with GLIM to achieve a higher signal to noise ratio (SNR) compared to widefield GLIM^32^. This method, called laser-scanning GLIM (LS-GLIM), maintains excellent depth sectioning because of the confocal operation and high numerical aperture (NA) optics and was shown to be effective with highly scattering samples^32^.

While being highly sensitive to the optical phase, QPI techniques are not highly specific to the structures/chemicals of interest in the sample. Deep learning advances are bringing effective solutions to address this shortcoming^33, 34^. Kandel et al. ^35^ combined deep learning with QPI to introduce computational specificity, and named the technique: Phase Imaging with Computational Specificity (PICS) ^35^. In PICS, deep learning models are trained on pairs of phase and fluorescence images (stained to detect structures of interest such as cell nuclei and cell cytoplasm). Post-training, the deep learning models are used to predict structure-specific fluorescence-labeling information from unstained phase images alone. Chen et al. ^32^ merged LS-GLIM with PICS in a technique called artificial confocal microscopy (ACM) to predict confocal quality fluorescence images from LS-GLIM phase images in 3D. This method thus enables label-free, quantitative phase tomographic imaging of highly scattering samples with computational specificity ^32^.

This paper presents a machine-learning-enabled health viability assay for mouse embryos using ACM. We first demonstrate the application of ACM in the label-free detection of nuclei in mouse embryos. By utilizing the optical phase information detected by LS-GLIM, we can extract insights about the structure and composition of the detected nuclei and the entire embryo which are then used to classify the health of an embryo into two classes: healthy/intermediate or sick. Furthermore, our imaging technique is sensitive to changes in protein concentration distribution through optical phase measurements. While the system does not aim to provide specific information about individual proteins, it can quantify changes in the distribution of the overall protein concentrations between the cells of the embryo. To our knowledge, this is the first health-grading study combining the embryo’s structural and compositional features from the same instrument in a label-free, non-invasive manner.

## 2. Results

### Workflow

The workflow for the study is shown in Fig. 1. A laser-scanning confocal microscope equipped with a gradient light interference microscopy (GLIM) module in the transmission path is shown in Fig. 1a and was used for embryo imaging in this study. This system produces a phase image (Fig. 1b) and a corresponding fluorescence image (Fig. 1c) for the same field of view.

**Figure 1.**
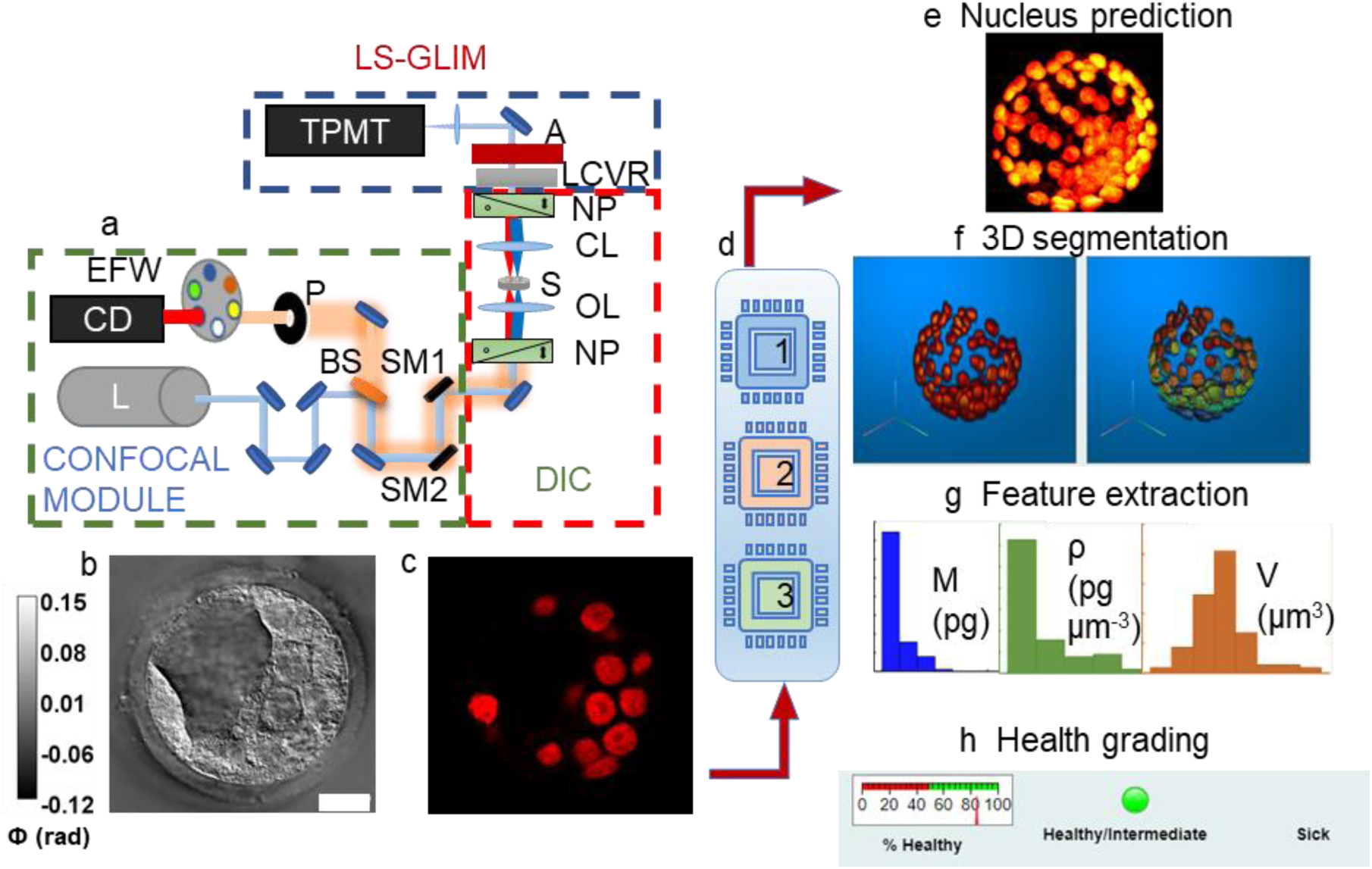
Workflow: a. ACM system setup, NP: Nomarski prism, OL: objective lens, CL: condenser lens, LCVR: liquid crystal variable retarder, BS: beamsplitter, P: pinhole, EFW: emission filter wheel, SM1/SM2: scanning mirrors, A: analyzer, S: sample, CD: confocal detector, L: laser source, blue dotted box depicts the LS_GLIM module, red dotted box depicts the DIC microscope without analyzer, and green dotted box depicts the confocal module. b. Final GLIM image c. corresponding fluorescence image stained for nuclei identification in the embryo slice shown in b. d. Deep learning modules: 1: Nucleus prediction model (NPM), 2: Feature-based health grading model (FBM), 3: Image-based health grading model (IBM) e. Composite of nucleus prediction, f. 3D rendering from 2D predictions and corresponding segmentation labels per nuclei, g. An example of extracted nuclei features (dry mass M, dry mass density ρ, volume V) h. Health grading of embryos by IBM and FBM. Scalebar 20 µm for b and c. Colorbar in b represents optical phase distribution (ϕ).

Using this system, we can extract dry mass and dry mass density (see Methods Section), which are intrinsic markers, from label-free phase images obtained using LS-GLIM. These quantities are linearly related to phase information of the sample, representing the non-aqueous content of the sample and hence relate to the protein/DNA concentrations in the biological samples^24^.

Three machine learning models were developed and trained (Fig. 1d). Model 1 is a nucleus prediction model that can identify nuclei from the GLIM images. A maximum intensity projection of an embryo’s predicted nuclear content is shown in Fig. 1e. These 2D predictions can then be stacked into a 3D structure, and after segmentation, unique color-coded labels are assigned to individual nuclei in 3D through custom MATLAB code (Fig. 1f). Following 3D segmentation, features for each nucleus in each cell within the embryo are extracted (Fig. 1g). Model 2 is a feature-based classifier for the health assessment of the embryo. This model accepts tabular data of features calculated from the 3D segmentation mentioned above and classifies the embryo’s health into one of the two classes H/I (healthy/intermediate) and S (sick) (Fig. 1h).

Model 3 is an advanced image-based classifier for the health prediction of embryos directly from 3-channel GLIM images of an embryo without requiring nucleus prediction or segmentation.

### Label-free nucleus prediction model (NPM)

We trained an EfficientNet B0^36^-encoded UNet^37, 38^ to identify nuclei from phase images of mouse embryos. Paired phase and fluorescence images for the same field of view were used as input and target ground truth (GT) for the model training (Figs. 2a and 2b). The nucleus prediction results are shown in Fig. 2c. These images are from an unseen test dataset and illustrate the predicted nuclei align precisely with the nuclei visualized in the embryo with fluorescence. Supplementary movie M1 shows a z-evolution of an overlay of GLIM images (grayscale) with nucleus predictions (red) for an embryo. The fluorescence (Fig. 2d) and nucleus predictions (Fig. 2e) are stacked into 3D reconstructions, with the 3D volumetric reconstruction shown in the Supplementary movie M2. The model achieved a peak signal to noise ratio (PSNR) 36.94, multi-scale structural similarity index (MS-SSIM) 0.94, and Pearson correlation coefficient (PCC) 0.81, respectively on the unseen hold-out test-dataset (Fig. S1d).

**Figure 2.**
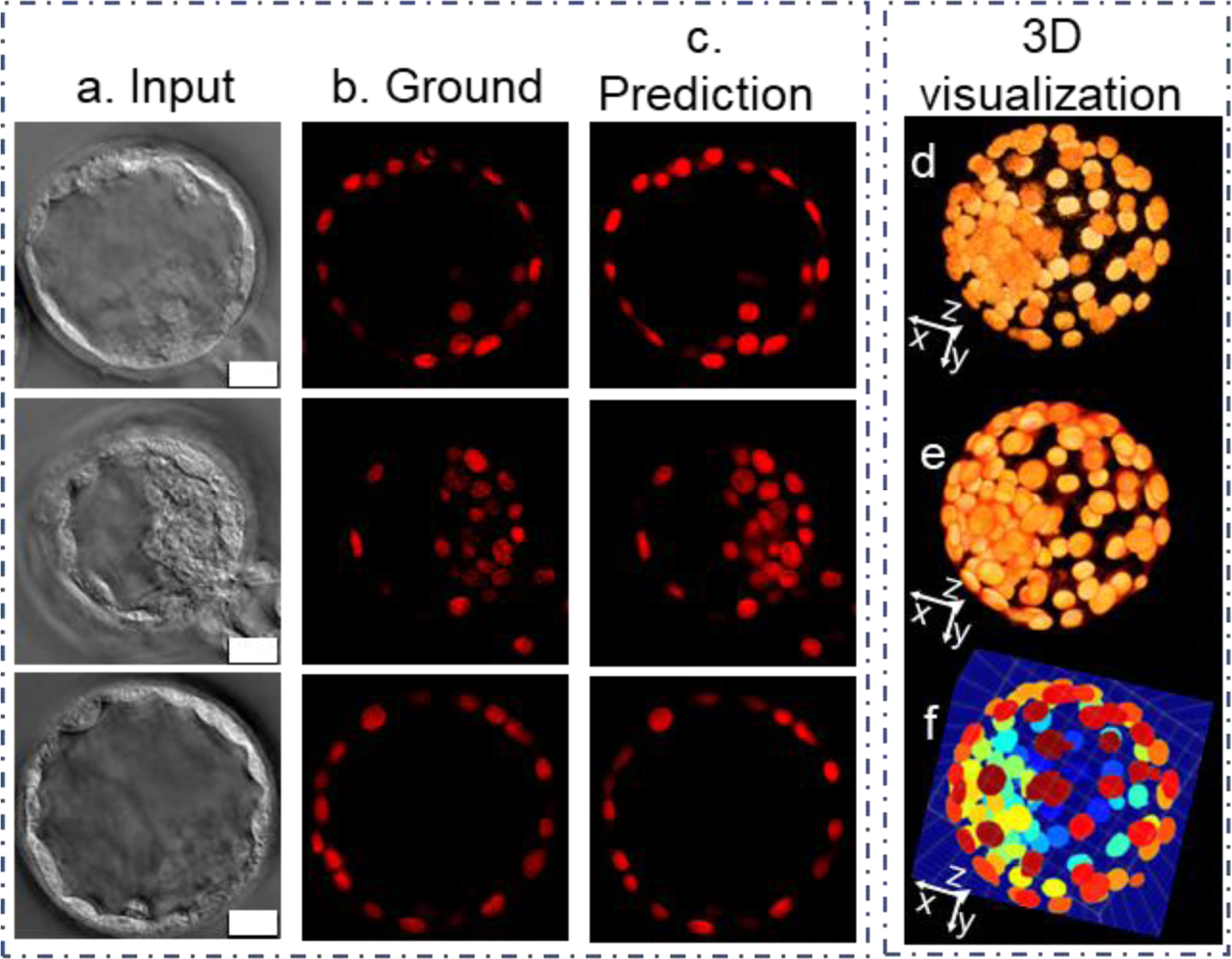
Nucleus prediction model results and 3D visualization: a. Input LS-GLIM z slice, b. Corresponding ground truth fluorescence image marking the nuclei, c. Model prediction. d. 3D stacked ground truth, e. 3d stacked prediction, f. 3d instance segmentation labels. Scalebar 20 µm for a, b and c.

### Nuclear feature extraction and insights

3D segmentation of the nuclear volume (Fig. 2e) gives an instance segmentation map (Fig. 2f). The labels are increasing in magnitude along the z-direction; therefore, a maximum value z-projection can serve as a nucleus count map providing critical information regarding the number of nucleated cells in each blastocyst (Fig. 3b, nucleus count map for the stacked nucleus prediction in Fig. 3a). After 3D segmentation, features were extracted at the level of both individual nuclei and the whole embryo. Three health classes of embryos were present in the dataset: healthy (H), intermediate (I), or sick (S), assigned by an embryologist. Histograms grouped according to the embryo’s assigned health class are shown in Figs. 3c-3i, with colors blue for healthy, green for intermediate, and red for sick embryos. Nucleus level features include dry mass (M (pg); Fig. 3c), dry mass-density (*ρ* (pg-*μ*m^−3^) ; Fig. 3d), surface area (S (*μ*m^2^) ; Fig. 3e), sphericity (Sp; Fig. 3f), and volume (V (*μ*m^3^); Fig. 3g). These distributions, along with the enclosed box plots, show these features differ between the healthy/intermediate (H/I) and the sick (S) classes of embryo. Sphericity and volume show a bimodal pattern for the S class indicating the presence of normal as well as fragmented nuclei. A total of 7788 nuclei from 152 embryos were analyzed, with health distributions according to class shown in Fig. 3j. All features assessed, with the exception of volume (Fig. ED1d), demonstrated statistically significant differences between the H/I and the S class (Table 3 and the boxen plots^39^ in Fig. ED1). Differences in the volume distribution were not statistically significant because of the variety of nuclear shapes found in the sick class that overlap with the healthy/intermediate class distribution (Fig. 3g and Fig. ED1). The H/I and S classes were also significantly different for the embryo-level feature ‘number of nuclei’ (Figs. 3h and ED1f). The sick embryos tended to have a slightly higher dry mass than healthy/intermediate embryos which can be attributed to the fact that most sick embryos had arrested development prior to blastocyst formation and were thus a ball of cellular material without cell differentiation into distinct trophectoderm (TE) and inner cell mass (ICM) cell populations and accumulation of blastocoel fluid. The changes in volume at the population level between late-stage morulae and blastocysts are insignificant. This explains the increase in dry mass density, which is a ratio of dry mass to volume. An enhanced level of nuclear degradation/fragmentation is also observed in sick embryos, which may explain the appearance of a bimodal distribution in sphericity and a large spread of volume distribution.

**Figure 3.**
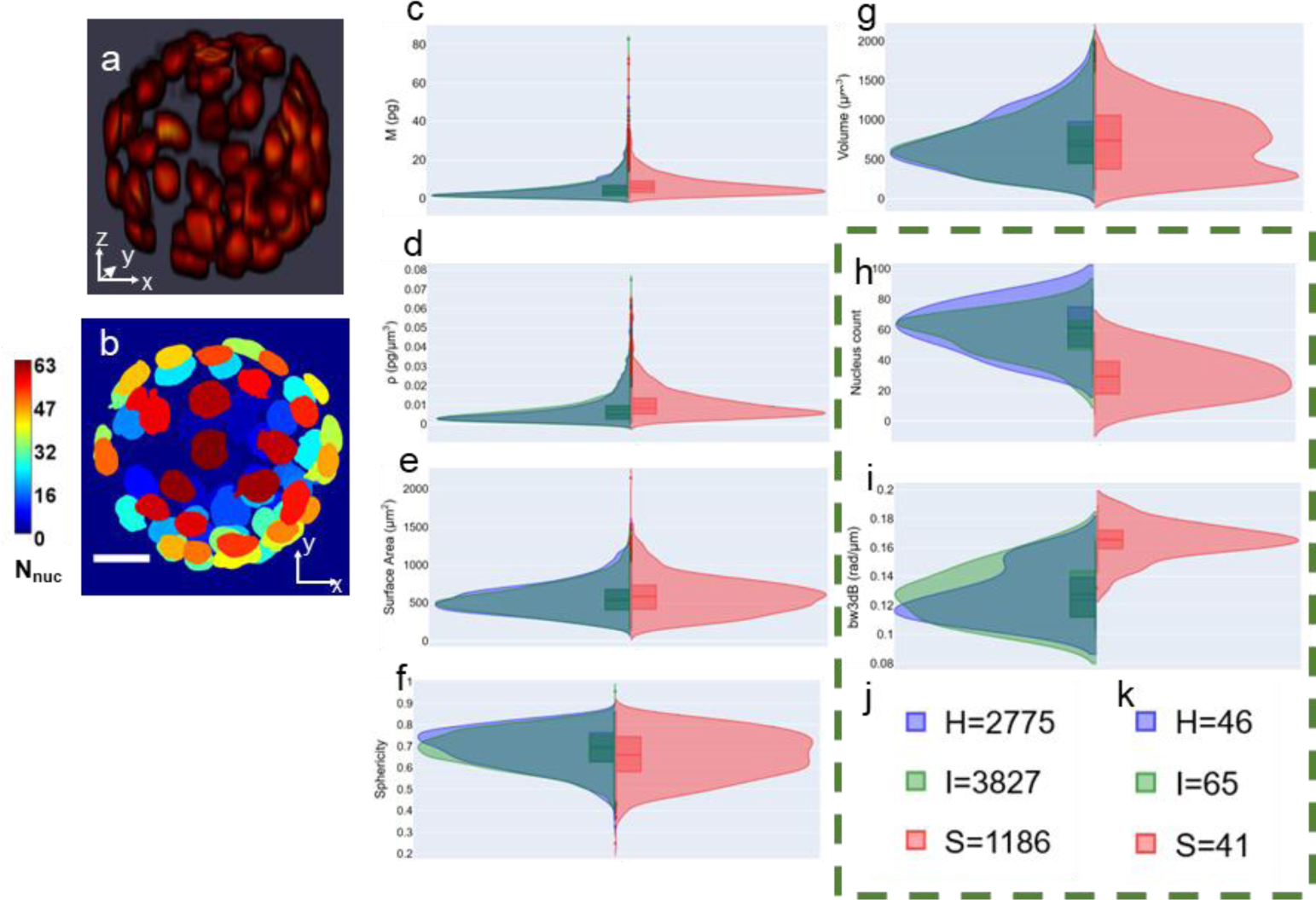
3D segmentation and related features: a. Stacked model predictions, b. maximum projection along z-axis of the labelled volume in a. showing the number of nuclei in the embryo. The colorbar shows the number of nuclei detected in the embryo, c-i. Histograms for nuclear dry mass, nuclear dry-mass density, nuclear surface area, nuclear sphericity, nuclear volume, embryo-wise nucleus count, and embryo-wise 3dB power bandwidth of scattering amplitude spectrum for 152 embryos grouped by health classes: Healthy (H) (blue), Intermediate (I)(green) and Sick (S)(red) class. For nuclei level parameters the number of nuclei in the three classes are shown in j and for the embryo level features the number of embryos per class are shown in k. Z-height of embryo in a and b is 91 µm. Scalebar for b is 20 µm.

Supplementary movie M3 shows 3D stacked GLIM images from the whole embryo, followed by the nucleus predictions, segmentation labels, and nuclear dry mass density distribution. The nuclear dry mass density distribution shows the mean dry mass density map per nucleus and implies that each nucleus represents the mean dry mass density averaged over its nuclear volume. The gradient in the colormap is due to Gaussian filtering applied to the 3D rendering.

### Nucleus arrangement and scattering amplitude spectrum: information in 3D arrangement

To better understand the implications of the 3D arrangement of nuclei within an embryo and the potential prediction of embryo health by this feature, we modeled an embryo as a scattering system of identical repeating units placed randomly in the spatial domain. We applied scattering theory to the remodeled nuclear distribution within an embryo and computed scattering amplitude and extracted the half-power bandwidth-bw3dB^24^ (see Methods Section). The bw3dB distribution is shown in Fig. 3i and was found to be statistically different between the H/I and S classes (Table 3).

For the embryo-level features, sick embryos had a lower nuclear count (Figs. 3h, ED1f and S4) and a larger scattering amplitude spectral bandwidth than H/I embryos (Figs 3i and ED1g). The maximum phase projections of the LS-GLIM images from a set of embryos belonging to the three classes: healthy (enclosed in a green box), intermediate (enclosed in a blue box), and sick (enclosed in a red box) are shown in Fig. S5a. Figures. S5b and S5c represent the maximum intensity projection of actual fluorescence and nucleus detection model predictions for the embryos shown in Fig. S5a, respectively. The maximum intensity projection of the scattering amplitude spectrum for the respective embryos is shown in Fig. S5d. While healthy or intermediate embryos had a well-defined and tightly focused power spectral density, sick embryos exhibited a diffused spectrum.

### Blastocyst stage embryos display distinct nuclear dry-mass density distribution between ICM and TE cells

Figure 4a shows the normalized nuclear dry-mass density map (see Methods Section) for all 152 embryos, with some selected examples shown in Fig. 4b. We observed that while cleavage stage embryos display no special mean dry mass density distinction between the nuclei (Fig. 4b, red boxes, sick embryos and first orange box, an intermediate embryo), blastocyst embryos (Fig. 4b, green boxes, healthy embryos) show a distinct pattern of nuclear dry-mass density distribution between TE cells and the ICM cells. It is readily observed that the TE cells have a higher mean nuclear dry mass density as compared to the nuclei of the ICM cells. As pointed out earlier in the manuscript, dry mass density is an indicator of DNA/protein content. Therefore, differences in the nuclear dry mass density suggests the DNA/protein concentrations differ in the nuclei of TE and ICM cells. Our results agree with reported differences in chromatin organization between the TE and ICM nuclei^40^. The pattern of differences in the nuclear dry mass density also hints at a correlation with the reported differences in the metabolic states of TE and ICM cells^21^.

**Figure 4.**
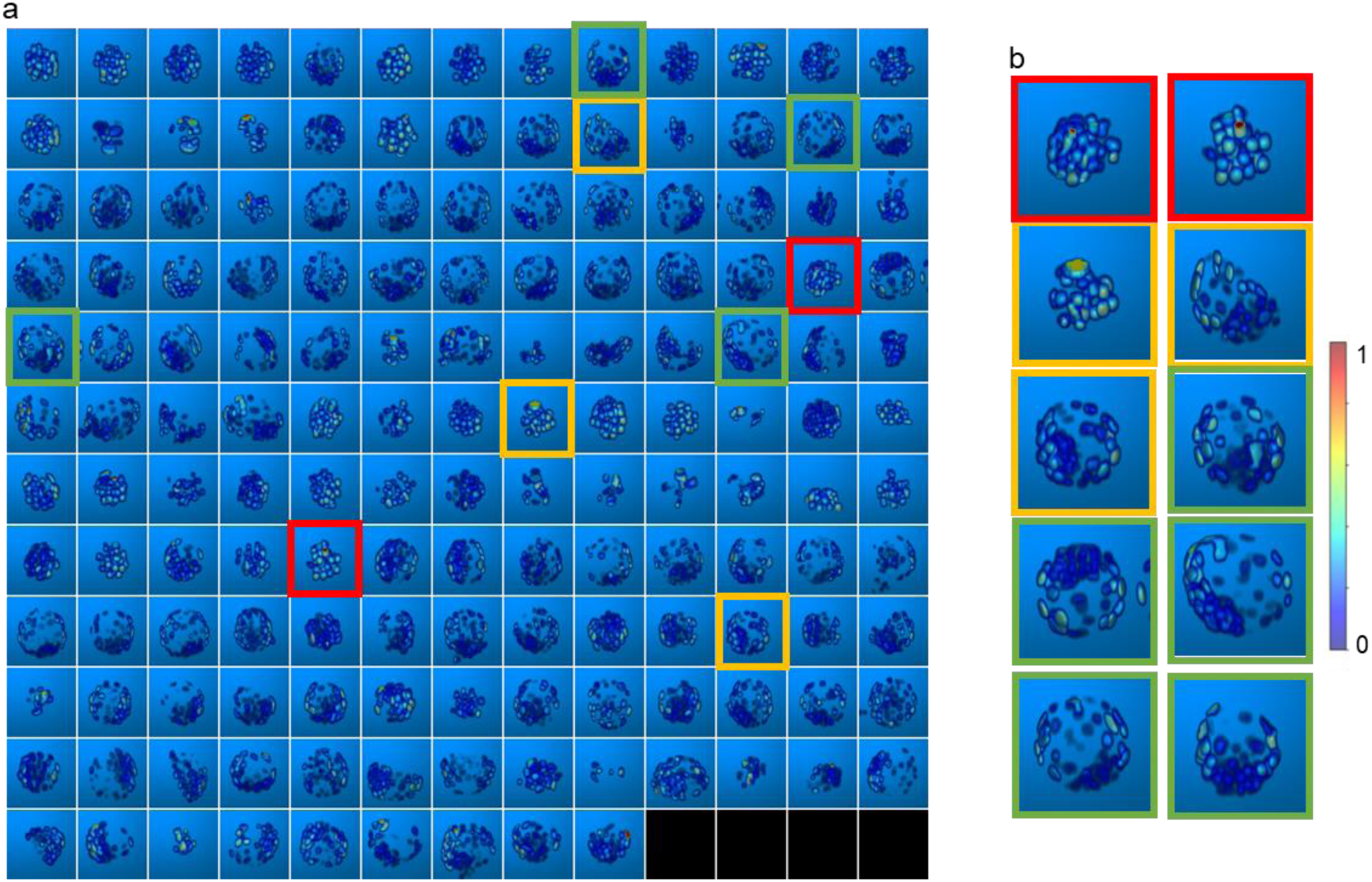
Mean nuclear dry-mass density distribution: a. 3D reconstructions of mean nuclear dry mass density map for all 152 embryos. b. Zoomed-in maps for selected embryos from a, enclosed by red boxes (sick), orange boxes (intermediate), and green boxes (healthy).

### Embryo viability assessment based on the extracted features and feature based health grading model (FBM)

After feature extraction and significance testing, features were analyzed for cross-correlations. The heatmap in Fig. ED1h depicts the correlation matrix. Based on the correlation and statistical significance (see Methods Section), five features that convey information about the structural and compositional aspects of the nuclei and embryo were selected for classifier training: nucleus shape (surface area and sphericity), nucleus organization in the embryo (bw3dB), number of nuclei in the embryo (nucleus count) and the mean protein/DNA content of each individual nucleus (dry mass density). A neural network classifier with two hidden layers was trained on the selected features for classifying individual nuclei to either the H/I or S class. We call this model a feature-based health grading model (FBM). The health of the whole embryo was then inferred by combining nuclear-based decisions per embryo in a max voting procedure as described in the Methods Section. The embryo-level confusion matrix and the performance metrics (precision, recall, and F1-score), evaluated on the blind-test dataset of 72 embryos (Supplementary Table1), are shown in Fig. 5a. Corresponding nucleus level results are shown in Fig. ED2a. This model achieved an F1 score of 1 for H/I and S classes at the embryo level health grading (Fig. 5a). At the nucleus level, the F1 scores were 0.99 and 0.94 for H/I and S classes, respectively (Fig. ED2a). The test set of embryos was imbalanced with respect to the proportion of nuclei and embryos in each health class. However, the significance of detecting healthy or intermediate embryos was considered a more critical task than detecting sick embryos, and we therefore weighted our metrics with the frequency of individual classes within the test dataset. The weighted F1 score for the classification model at the embryo level is 1, and at the individual nucleus level is 0.98. As the results indicate, this model could classify the health grade of a nucleus belonging to an embryo with an AUROC 0.986±0.007 (Fig. ED2a) for S class embryos.

**Figure 5.**
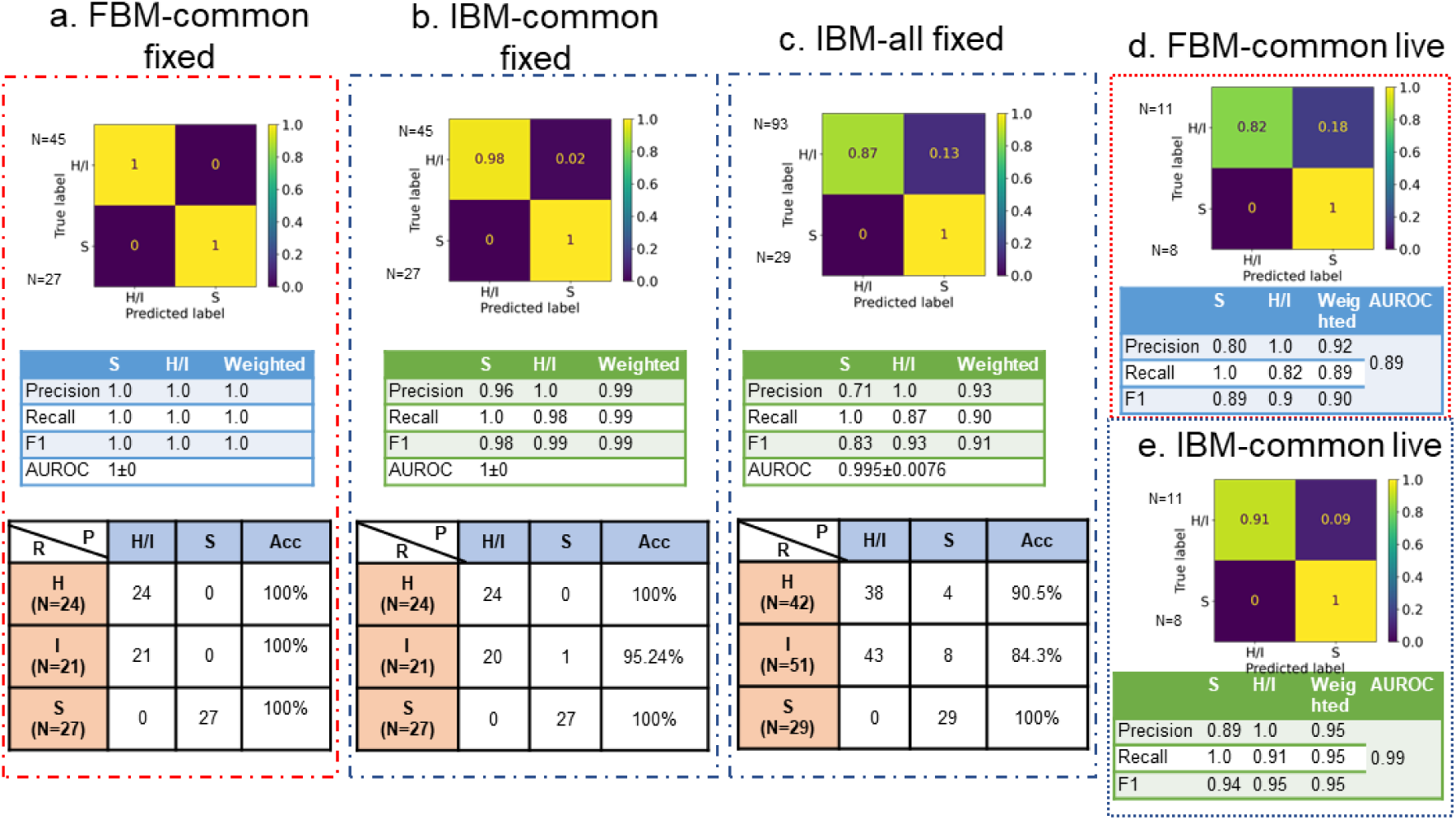
Health grading model performance-confusion matrix, performance metrics and real-class wise performance for: a FBM for common test set of 72 embryos b. IBM for common test set of 72 embryos c. IBM for extended data of 122 embryos d. FBM for live embryos (19 instances) and e. IBM for live embryos (19 instances). R denotes the real class by expert (healthy (H), intermediate (I) and sick (S)) and P denotes the model predicted class (H/I and S), N denotes the number of embryos per class, Acc denotes the accuracy per real class.

### Health grading directly from LS-GLIM images and the image-based health grading model (IBM)

We trained a second model, the image-based health grading model (IBM) to enable health grading directly from LS-GLIM images. This model was a pretrained EfficientNet B7^36^ classifier trained in a transfer learning approach with some architecture modifications (see Methods Section). The input to the model was a three-channel GLIM image, hereafter referred to as a z-slice, where each channel represents a consecutive z-section spaced 1 µm apart. This type of input was chosen to provide the model with a correlative view/relationship between the neighboring z-sections. The model was evaluated on individual z-slices. For the prediction of the embryo’s overall health, predictions from individual z-slices were combined by max voting (described in the Methods Section). The resultant confusion matrix is shown in Fig. 5b for the same test set of 72 embryos used to evaluate FBM in Fig. 5a. IBM achieved an approximately equal performance as the FBM, with embryo-level F1 scores for H/I and S classes being 0.99 and 0.98, respectively (Fig. 5b). The classifier’s weighted precision, recall, and F1 score for individual z-slices were 0.92, 0.91, and 0.91, respectively (Fig. ED2b), with 0.981±0.0015 AUROC. Of note, the max voting removed noisy predictions making it more accurate than the z-slice level predictions.

We tested the IBM on an expanded test dataset of 122 embryos (that included samples used in the training set for the FBM but unused for the IBM combined with 72 embryos from common test set) to explore the cause of one misclassification in the common test dataset. The IBM still performed well with a weighted F1 score of 0.91 at the embryo level (Fig. 5c). The z-slice level results are shown in Fig. ED2c, where the model achieved a weighed F1 score of 0.88. The classification matrix (bottom row) in Fig. 5c, indicates that most of the misclassifications involve intermediate (I) embryos, that were misclassified as S class embryos by the model. It is essential to note that neither the FBM nor the IBM misclassified an S class embryo as H/I class.

### Model performance on live embryos

All previous analyses were performed using fixed mouse embryos. To evaluate the application of the proposed embryo assessment tool in a clinical setting, a second test set of 8 live embryos was analyzed providing a new dataset of 19 time-instances from 8 live embryos (11 instances of embryos expert embryologists marked as H/I class + 8 marked as S class, Supplementary table 2). This represented an out-of-distribution dataset because training of the models was performed on fixed embryos. Figures 5d, and 5e show the confusion matrix for the FBM and IBM evaluated on this test set, respectively. As above, these results are the base models’ embryo-level predictions obtained after max voting. The IBM outperformed the FBM on live embryo health classification with a weighted F1 score of 0.95 compared to 0.90 (Fig. 5e and 5d, respectively). The visual results from individual examples of embryos from the two test sets are shown in Fig. 6. Figures 6a, b show three examples of test set-1(fixed embryos). Figures 6c, d display the 24-hour time-lapse of one embryo. Two different live embryos are shown in Figs. 6e,f. We observed that the IBM misclassified an intermediate embryo (middle embryo in Fig. 6a) as S class in the fixed embryo dataset and again one intermediate embryo as S class in live embryo dataset (Fig. 6e), and the FBM misclassified two healthy live embryos as S class (Figs. 6c-6f). We noted that this misclassification pattern was observed in most of the errors made by the IBM and FBM. As a result, some intermediate embryos were assigned to the S class. This small proportion of “down grading” misclassifications (Fig. 5) was in accordance with our proposed aim stated earlier that puts a greater emphasis on the accurate detection of healthy embryos.

**Figure 6.**
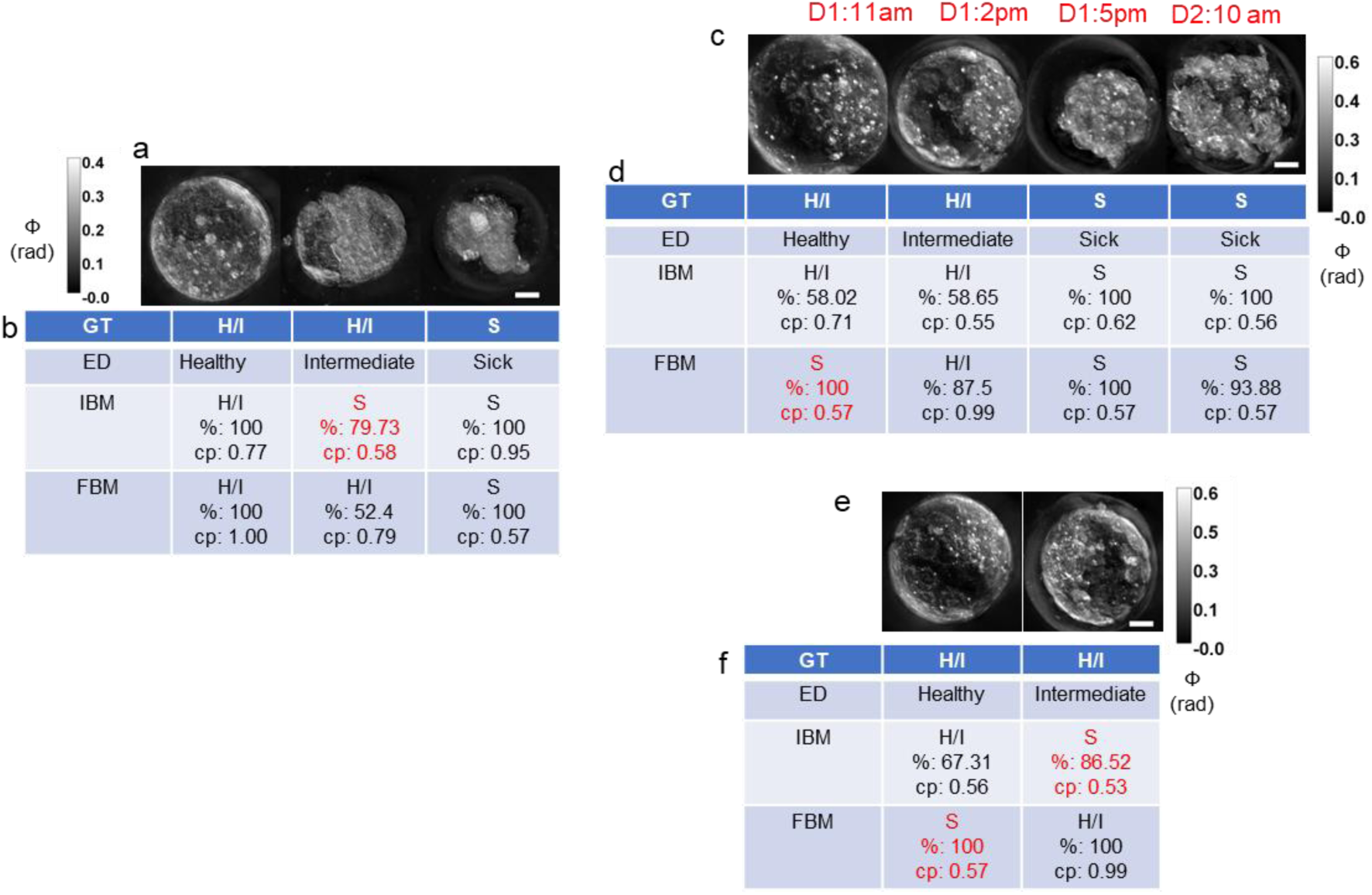
Health grading of embryos: a. LS-GLIM composite images from common test set of fixed embryos b. Model predictions on test embryos in a. c. Time-lapse LS-GLIM composite images from common test set of live embryos d. Model predictions on test embryo in c. e. LS-GLIM composite images from common test set of live embryos showing two different embryos f. Model predictions on test embryos in e. Red entries in b, d and f represent wrong predictions. Colorbars show phase distribution (ϕ). Scalebar is 20 µm for all images. GT: ground truth class, ED: real class assigned by experts, IBM: image-based classification model, FBM: feature-based classification model. H/I: healthy or intermediate class, S: sick class. cp: average prediction score over majority predictions, % denotes percentage of majority predictions (z-slices for IBM and nuclei for FBM). D1, D2: Days of time-lapse followed by timestamps of acquisition.

We believe that the decrease in the performance of the FBM in the case of live embryos is because of the nature of the damage to the embryos. While the training data contained sick embryos that were growth-halted naturally or by treatment (see Methods Section), the live embryo test data was obtained from blastocyst embryos that may have undergone degradation over time. If the nuclear degradation was determined by the model to be high, the FBM classified the embryo into S class. The change in phase distribution of the live samples (Fig. ED3a) may also be a contributing factor as it affected the dry mass density distribution. Changes in the surrounding media of the samples impact the phase distribution profile^41^. Furthermore, fixation also causes cell shrinkages and altered refractive index^42^. These differences in cellular integrity result in a shift in data distribution in the live embryos compared to the training data on fixed embryos. Hence, we refer to the data from the live embryos as an out-of-distribution dataset. Overall, the performance of the FBM is acceptable (weighted F1 of 0.9, Fig. 5d) under the above-mentioned distribution shift.

Data distribution shift did not appear to impact the IBM since changes in the phase values are minimized by image normalization during preprocessing. In addition, the IBM looks at the whole embryo rather than just the nuclei and therefore has a much larger feature space than the FBM, which may explain its superior performance over the FBM for live embryos.

### Understanding the models: Interpretation of FBM and IBM

We deployed model interpretation methods to understand the model decisions. For the FBM, we used Shapley additive explanations (SHAP) ^43, 44^ calculated in Python 3.9.7 using the SHAP API over the correct test (in-distribution test set) observations to indicate feature importance (Figs. ED3b-c). The feature importance bar graph in Fig. ED3b shows the FBM pays greatest attention to the nucleus count (*nuc_count)* and the scattering amplitude spectrum 3dB bandwidth (*bw3dB),* highlighting nucleus distribution inside the embryo volume. The third most important feature is the sphericity of the nuclei (*sphericity)* followed by compositional information from dry mass density distribution (*dmd)*. *surface area* is the least important feature, which was expected because of the lower statistical significance as compared to the other parameters (Supplementary Table 3). These results indicate that the FBM focuses more on the embryo size and nuclear density and distribution, and further explain the lower performance of the FBM on live embryos, as hypothesized above. The degenerative changes seen in live embryos differ from those observed in the growth-arrested embryos in the training dataset, such that the live embryos may have undergone degradation after reaching the blastocyst stage. However, from the feature importance plots (Fig. ED3b-c), we can infer that the change in dry mass density values is less detrimental to the model performance than the changes in the degradation mode (thus affecting nucleus number/distribution). We also determined the effect of feature values on the FBM predictions. The SHAP summary plot in Fig. ED3c shows that a low nuclear count, high scattering amplitude spectrum bandwidth, low sphericity of nuclei, and a high nuclear dry mass density favored a prediction of S class. These results matched our observation of individual histograms of features (Figs.3 and ED1). However, the surface area feature reported high values in both health classes and did not match the histogram explanations of high surface area being associated with S class nuclei.

For the IBM interpretation, we used the GradCAM approach^45, 46^. For the correct predictions of each class, a blind test was performed between the experts and the model GradCAM to highlight the regions of importance in each image relevant to the health grade of the embryo. We used randomly selected z-slices from the test set embryos for this study. Extended data figure ED4 shows the cases where the GradCAM matched well with the expert markings for healthy (Fig. ED4a-e) and sick (Fig. ED4f-j) embryos. With H/I class embryos, the model seemed to focus on TE cells (Figs. ED4 a, b and d) and the ICM (Fig. ED4 c and e). In contrast for the S class embryos, the model focused on abnormal/individual blastomeres (Fig. ED4f, g, i) or cytoplasmic fragments (Fig. ED4 f-j). Overall, in S class embryos the model focused on features that matched the annotated regions marked by embryology experts. However, there were some H/I embryos where the expert annotations and the model GradCAMs did not match perfectly (Fig. ED5). In Fig. ED5a, in addition to the region marked by the experts, the model was also focusing on the blastocoel-like cavity. In Fig. ED5b and d, the region of interest was out of focus in the current z-slice and the model appeared to ignore such out-of-focus features. Figure ED5c shows another example of mismatch, where the cells had aberrant structure, which may explain why the model ignored the marked region. Finally, there existed cases like Fig. ED5e where the model may pay attention to specific features inside the cell cluster.

We also explored cases where our model produced incorrect results (Fig. ED6). We included two random z-slices per embryo where the predictions were incorrect at the slice level. Figures ED6a-d show two z-slices per embryo of H/I embryos predicted as S class at the z-slice level. We suspect that the model focused on the abnormal shapes of the embryos at these z-slices. The bottom panel (Fig. ED6e-h) provides examples of z-slices of two S class embryos predicted as H/I at the z-slice level. It is important to mention that, except for the embryo in Fig. ED6a-b, all the wrong predictions at the z-slice level did not change the model-assigned grading at the embryo level as the misassigned z-slices were in the minority voting category. However, the embryo in Fig. ED6a-b was predicted as S class at the embryo level, meaning that the model marked the majority of the z-slices of this embryo as S.

### Sparse z-slice predictions of IBM

This is a preclinical study to demonstrate a method that can be deployed in the clinical setting for real-time health grading of human embryos. ACM requires very low illumination power compared to fluorescence-based microscopy techniques. However, scanning through a whole embryo (of ∼100 µm diameter) can take approximately 40-50 minutes. To minimize the scan time and further reduce exposure of the embryo to light, we tested the IBM on a few z-slices instead of the whole embryo z-scan. The test was performed on the live embryo test dataset. It is important to note that one z-slice comprises a 3-channel image of three consecutive 1µm spaced z-sections that were acquired irrespective of the inter-slice distance or the number of z-slices. We removed intermediate z-slices in the already-acquired dataset to mimic the increase in z-step size from 1 µm to 60 µm in steps of 5 µm. Figure ED7 shows results from the test on 18 live instances of embryos where the IBM predicted correct results. These data indicated that the maximum inter z-slice distance below and at which the correct predictions are maintained is 10 µm creating an allowable z-steps range for the z-scan of an embryo using a 40x/1.3 NA objective to be 1 µm-10 µm. We also investigated the minimum number of z-slices required to maintain the correct prediction (Fig. ED8). Since the predictions within 1 to 10 µm are correct, we can see that the number of z-slices for this range of interslice distance (1-10 µm) is greater than 5. An effective threshold minimum number of z-slices was determined to be 7. We avoided using 6 z-slices to prevent the occurrence of an equal percentage of predictions for both H/I and S classes that would yield an inconclusive result. Acquiring 7 z-slices with an interslice distance ranging from 1 µm-10 µm will lead to ∼80-85% reduction in scan time per embryo.

## 3. Summary and Discussion

Label-free estimation of nuclear structures and the assessment of the viability of embryos is a new direction of research in embryology. In this work, we presented a label-free computational imaging technique to perform the following tasks: 1) detect the nuclei in embryos using NPM, 2) obtain information about structure and composition from the same instrument in a non-invasive, label-free manner, 3) segment and analyze the nuclear properties, 4) associate these properties to the health grade of the embryo. We also devised a new biomarker (scattering amplitude spectrum bandwidth) based on the scattering theory to use as a predictor of the health grade of the embryo. In addition, intrinsic markers (dry mass density) and shape descriptors (nuclear surface area and nuclear sphericity), along with the total nucleus count in an embryo, were found to be effective predictors of embryo health. Indeed, the nucleus detection task is of paramount importance as it conveys information regarding intracellular well-being and appropriate embryo development that is not discernable from its external morphology^47^. Our results also provided new data regarding differences in nuclear dry mass density between TE and ICM cells.

We trained two types of embryo viability assessment models: the FBM and the IBM. Both models performed equally well on the in-distribution blind test dataset of fixed mouse embryos. Most instances (8 out of 12) of misclassification by the IBM involved intermediate embryos that were classified as sick. This model approach aligns with our philosophy behind the proposed viability assessment which places more emphasis on the accurate detection of healthy embryos.

When the two models were tested on an out-of-distribution (live embryo) dataset, the FBM suffered an ∼11% decrease in AUROC compared to its performance on the fixed embryo dataset, while the IBM performed approximately the same (0.5% decrease in AUROC). We attribute the decrease in FBM performance to the data-distribution shift in the live embryos. However, the FBM performed acceptably (weighted F1 of 0.90), even after the distribution shift which represented a baseline model performance. We also tested the IBM for cases of sparse predictions, and the results indicate that it can accurately grade embryo health from 7 z-slices per embryo spaced up to 10 µm apart reducing the total time required for assessment of each embryo.

A key strength based on an assessment of 122 embryos for IBM and 72 embryos for FBM is that both models never assigned an incorrect class to a S class embryo and would therefore permit effective removal of non-viable embryos from a clinical cohort. We also demonstrated that our models track the health of the embryos in a time-lapse acquisition. This has potential implications for incorporation of the GLIM module into a standard TLS system.

The work by Khosravi et al. ^17^ is one of the most relevant studies in this area of embryo health assessment using machine learning. They used a large dataset of multi-focal images and time-lapse imaging data to develop a deep learning framework (“STORK”) for grading embryos into two classes: good or poor. They reported an impressive AUC of 0.987 for the in-distribution dataset and an AUC of 0.90 and 0.76 for out of distribution dataset, using images from different clinics. However, ‘fair’ (equivalent to our intermediate class) quality embryo images were discarded from the training dataset. In our method, we did not exclude ‘intermediate’ embryos from our training dataset. Our embryo viability assessment method is highly accurate with an AUROC: (0.995, 1) and (0.99, 0.89) by (IBM, FBM) for the in-distribution and out-of-distribution datasets respectively. Our image-based model thus outperforms the previous study^17^ in terms of generalizability to the out-of-distribution dataset which is likely attributed to the highly sensitive quantitative phase information in our images provided by LS-GLIM and the completeness and diversity of the training dataset because of the inclusion of all three types of embryos: healthy, intermediate, and sick. In addition, our work also included insights into embryo morphology and composition utilizing phase information.

Other reports document studies with blastocyst grading algorithms incorporating ground truth data of known pregnancy outcomes, either through fetal heartbeat^48^, PGT-A^49^, or beta-hCG^49, 50^ based pregnancy results. The Life Whisperer AI model by VerMilyea et al. ^48^ (accuracy: 64.3%), Embryo Ranking Intelligent Classification Algorithm (ERICA) by Chavez-Badiola et al. ^49^ (accuracy: 70%), EmbryoNeXt by Marsh et al. ^51^ (AUC: 0.869 and 0.807 at 2 and 3 hours after thawing the embryos, respectively), the study by Tokuoka et al. ^12^ (accuracy: 83.87%) and the work by Berntsen et al. ^52^ (AUC: 0.95 when evaluated on the entire dataset and 0.67 when evaluated on embryos with known implantation data (KID)) are some of the similar studies for embryo viability assessments, birth outcome predictions or grading. Majority of these studies report either an image-based or feature-based classifier for embryo grading. However, in our work, we have demonstrated both types of models (IBM and FBM), and have directly compared their performances on the same embryo datasets. While the performance metrics of the models described in this study are higher than most of the other published studies, a direct comparison is not appropriate due to the different end goal and ground truth information, which is to grade the health of the embryo to determine suitability for selection for transfer, rather than determining an end point of pregnancy, with the ground truth being the health-gradings by the embryologists.

Despite the advantages mentioned above, our study also has some limitations. The first limitation is the drop in performance of the FBM due to the data distribution shift in live embryo culture conditions compared to the fixed embryos. This occurred due to an inherent problem of machine-learning models when presented with a shift in the data distribution. The FBM results represent a lower limit of the model performance in the case of a distribution shift. The FBM performance may likely be improved by including a more extensive and diverse training dataset with varying levels/modes of embryo/cellular degradation and sample preparation (fixed and live). A second limitation is the use of mouse embryos. However, the health grading tool we report herein can be translated to human embryos in the clinic with further training in future studies. The third limitation is that we did not compare the health of the embryos to pregnancy outcomes. This is because of the experiment design, where our training dataset was composed of fixed embryos. Additional studies are needed to combine our model’s health prediction with pregnancy outcomes. A second top-up model can be trained to map our model’s health grading on live embryos to pregnancy outcomes. Lastly, the size of our dataset (195 fixed embryos) is small compared to the studies involving human embryos. All the data collected in this study was self-acquired using LS-GLIM rather than accessing data from multiple sources (clinics).

In summary, our technique lays the foundation for a complete evaluation of embryos from the structural and compositional point-of-view without adding external reagents or invasive measurement procedures. Our method significantly exceeds the range of information provided by a standard TLS system regarding the 3D quantitative insights of the embryo^17^. Furthermore, our method provides flexibility to researchers/clinicians by using two types of models: a quick IBM model that is capable of sparse z-slice-based predictions (suitable for busy clinics or sensitive embryos) and an in-depth combination of NPM and the FBM (suitable for researchers) that provides a complete 3D quantitative assessment of the embryos.

## 4. Methods

### Embryo culture and staining procedure

#### Animals

Sexually mature B6D2F1 males and CD1 females at 35-42 days old were purchased from Charles River Laboratories (Wilmington, MA). The mice were housed at the Carl R. Woese Institute for Genomic Biology at the University of Illinois, Urbana-Champaign. All the mice were provided with feed and water ad-libitum and housed in individually ventilated cages under a controlled environment. Animal rooms were maintained at 22 ± 1 ℃ with a 12-hour light cycle. B6D2F1 males were housed individually for breeding setup purposes. Animal handling and procedures were performed in accordance with the University of Illinois Institutional Animal Care and Use Committee.

#### Chemicals

Ovarian superovulation was achieved using pregnant mare serum gonadotropin (PMSG, Cat. Nor-272-a, Prospec, East Brunswick, NJ), and human chorionic gonadotropin (HCG, Cat. C1063-1VL, MilliporeSigma, Burlington, MA). Materials for in-vitro culture, including potassium simplex optimized medium (KSOM, Cat. MR-121-D) and hyaluronidase, were purchased from MilliporeSigma (Burlington, MA) and sterile mineral oil (Cat. ART-4008-5P) from CooperSurgical (Trumbull, CT). OMOPS handling medium consisted of MgSO4 (1.2 mM); Glucose (0.5 mM); L-Lactate (6.0 mM); GlutaGRO (1.0 mM); Tarine (0.1 mM); NEAA (1x); EDTA (0.01 mM); Alpha Lipoic Acid (10 uM); Gentamicin (10 ug/ml); Hyaluronan (0.125 mg/mL); NaHCO3 (5.0 mM); MOPS (20.0 mM); pyruvate (0.2 mM); Citrate (0.5 mM); FAF BSA (4.0 mg/ml). A pure phthalate mixture was made by calculating and combining the appropriate amount of each phthalate according to the following percentages 35% DEP, 21% DEHP, 15% DBP, 15% DiNP, 8% DiBP, and 5% BBzP in 0.05% DMSO. Phthalates were purchased from Sigma. In previous, preliminary studies, incubation of mouse embryos in this phthalate mixture, at 1 µg/mL, reduced blastocyst formation by 50%. This concentration was used to produce the sick batch of embryos.

#### Embryo collection and in vitro culture

To induce superovulation, 35 to 42 days old female mice were injected intraperitoneally with 6 IU of PMSG, followed by 6 IU of HCG 48 hours post PMSG. Injections were performed at 2:00 to 3:00 pm. After the HCG injection, female mice were housed individually with male mice overnight. The following day, at approximately 20 hours post-hCG, the female mice were inspected for a copulation plug and euthanized by CO_2_ inhalation and cervical dislocation. Oviducts were collected aseptically and placed in a pre-warmed OMOPS medium supplemented with 5% fetal bovine serum (FBS, Atlanta Biologicals) and were maintained at 37℃ during collection and transportation. Embryos were released from the oviducts into the OMOPS medium using forceps to gently tear open the ampulla. The embryos were rinsed a minimum of three times with OMOPS medium before being introduced into a droplet of OMOPS medium containing hyaluronic acid (500 ug/ml) for cumulus cell removal. Exposure to hyaluronic acid was limited to a maximum of one minute. The embryos were rinsed two more times in clean OMOPS droplets until cumulus cells were removed. Cumulus-free embryos were placed into pre-equilibrated KSOM droplets under mineral oil. All the embryos were cultured in a ThermoFisher 8000 WJ CO_2_ incubator at 37 ℃, 6% CO_2_, and 80% relative humidity for five days at which point they were assessed using an Olympus IX70 phase contrast microscope.

### Fixation of Embryos and Immunofluorescence Staining for 7-AAD

The embryos were fixed in freshly prepared 50:50 methanol:acetone at 1:1 volume ratio at −20°C for 20 minutes. They were then transferred to Holding Medium (50 ml PBS + 0.25 g BSA (0.5%)), droplets covered with mineral oil, and maintained at 4°C until staining.

Immunofluorescence staining for 7-Aminoactinomcyin D (7-AAD) was carried out as follows. Fixed embryos were placed in permeabilization buffer (50 ml PBS + 500 µl Triton X-100 (1%)) for one hour. Embryos were then washed three times for 10 minutes in Washing Buffer (500 ml PBS + 500 µl Triton X-100 (0.1%) + 0.5 g poly-vinyl-pyrrolidone (0.1%)).

Embryos were then incubated with the 7-AAD Red Fluorescent Live/Dead Stain from Immunochemistry Technologies (#6163) at a 1:200 dilution for 30 minutes in the dark at room temperature followed by three 10-minute washes in Washing Buffer. Finally, embryos were mounted in 3 mm glass bottom dishes with 14 mm micro-well in 20 µl of mounting medium containing DAPI for 30 minutes in the dark at room temperature. Embryo droplets were covered with 100 µl of mineral oil, coverslipped and kept in the dark at 4°C for 24-48 hrs before imaging.

### Image acquisition and reconstruction

Embryos were imaged using a laser-scanning GLIM setup with confocal fluorescence detection for nucleus ground truth data^32^. A standard Zeiss Axio Observer microscope was equipped with an Airyscan LSM 900 module for fluorescence confocal operation and a GLIM module for quantitative phase detection for the same field of view. The system has two laser sources with wavelengths of 488 nm, and 561 nm. The microscope was operated using Zeiss Zen software for setting the experiment parameters, and the image acquisition was controlled by a custom build MATLAB code developed by Chen et al. ^32^. However, we modified the acquisition code for automated stage movement, coordinate tracking, and tomographic acquisition for multiple fields of view. All images were obtained using a Plan-Apochromat 40x/1.3 Oil DIC (UV) VIS-IR M27 objective with the pinhole set to 1.09 AU. The wavelength of the laser source for LS-GLIM was 488 nm, operated at 1% power with a detector gain of 350 and digital gain of 1. Four intensity images corresponding to phase shifts of 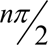, for n=0,1,2,3, were recorded at the transmission photomultiplier tube (T-PMT). From the 4-intensity images, a phase gradient image was extracted using the phase-shifting interferometric reconstruction algorithm as detailed in Nguyen et al. ^30^. The final phase image was obtained after applying Hilbert transform-based integration to the phase gradient image^30^.

Phase images convey information about the sample’s composition in terms of dry mass and dry mass density^24^ because of the linear relationship between dry mass density and phase. The relation between dry mass and phase can be expressed as:

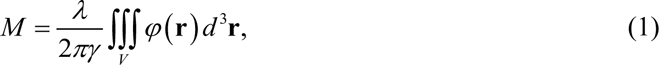

where, M is the dry mass, *λ* is the illumination wavelength, *γ* is the refractive index increment, and *φ* (**r**) is the measured 3D phase. The dry mass density is then evaluated as the average dry mass over the volume (V) of the sample.

For fluorescence detection of the 7-AAD nucleus signal (peak excitation and emission at 549/648 nm), excitation at 561 nm with 2% laser power, detector gain 750-800, and digital gain 1 were used.

The scan mode was set to frame for each channel, and pixel dwell time was 1.21 µs. In addition, the laser scan speed was set to 6 in a bidirectional scan mode. Tomographic acquisition of each embryo was done with a step size of 1 µm in the z-direction.

### Nucleus prediction model architecture and training details

The nucleus prediction model (NPM) is a UNet-style model with an Efficient Net-B0 encoder^36–38^. The architecture is shown in Fig. S1a with the submodule information in Fig. S1b. We first trained the model on a pair of images from LS-GLIM with corresponding images of the fluorescence-stained nuclei (Figs. ED9a and ED9b, respectively, from an unseen test dataset). The input image and the target image size were 1280 x 1280 pixels each. The fluorescence target image (Fig. ED9e) was median filtered with a window size of 5 to reduce spurious pixel noise (Fig. ED9f). The model predictions on the test set are shown in Fig. ED9c. The predictions also contain spurious background detections in the cell cytoplasm area because of the presence of a weak background signal in the fluorescence ground truth itself, more evident in Fig. ED9 b, c middle row (white arrows). To remove this background noise and increase the specificity of detection a preprocessing step was added for the target image. The histograms of the denoised fluorescence image were matched with the corresponding LS-GLIM image using the MATLAB function “*imhistmatch”*. The overall effect of this step was similar to thresholding, where negative pixels in the LS-GLIM (that mostly correspond to the background/cytoplasm) were zeroed out in the target fluorescence image. Another effect was that the target fluorescence image was much more specific now, such that slightly defocused nuclei and cytoplasmic signals were removed, as evident in Fig. ED9g. With the training set now containing images like Fig. ED9d as input and Fig. ED9g as the target, the model was able to remove spurious detections, as shown in Fig. ED9h. Images from the test set showing the predictions of both the models is shown in Figs. ED9i-ED9l, where Fig. ED9i is the input GLIM image, Fig. ED9j is the corresponding denoised target fluorescence image, Fig. ED9k is the first model prediction (without histogram matching), where the spurious cytoplasmic signals are also present (as indicated by white arrows) and Fig. ED9l shows the final model prediction (trained on a histogram-matched target), where the increased specificity is evident. The numerical results of both model comparisons are shown below each prediction. The final model shows an increased PSNR and MS-SSIM. The Pearson correlation coefficient (PCC) decreased after histogram matching. The visual results indicate the superiority of our final model. Thus, PCC was not found to be a good metric for this task.

The model was trained on a combination loss which is a linear combination of L1 loss, MS-SSIM loss, and Pearson loss defined as:

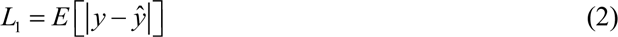

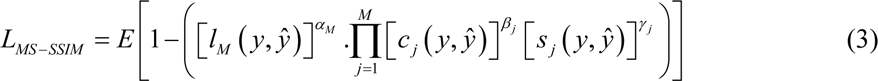

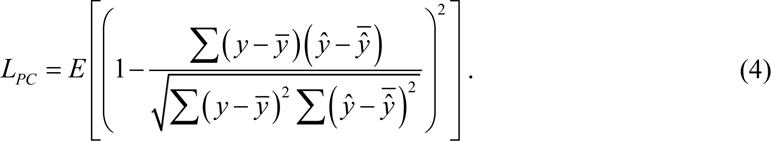

where, *l*, *c*, and *s*, are the luminance, contrast, and structure comparison measures as defined elsewhere^53^. MS-SSIM definition is followed from Ref. ^53^.

The final loss is

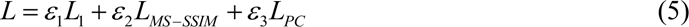

The weights of the loss components were determined empirically to be [2,1,0.5] for L1, MS-SSIM, and Pearson loss, respectively. Our choice of the loss function was inspired by a previous study^54^, where the authors have demonstrated the superiority of the L1 and MS-SSIM loss combination. The Pearson loss was used to overcome the image artifacts observed in previous studies (unpublished).

A mixed strategy^55^ of learning-rate warmup and cosine decay was followed for the learning rate, increasing it from 0 to the specified 1e-4 in the first 6 epochs and later decreased following cosine learning decay. From the loss curve (Fig. S1c), training was stopped after the 10^th^ epoch because no further improvement in validation loss was observed after the 8^th^ epoch. The low number of epochs is justified because of the large dataset employed for the training. Specifically, 4265, 1280 x 1280 pixels sized images were used for training that increased in number to 8530 training instances after augmentation. An additional 1434 images were used as the validation set, and a hold-out test set containing 1407 images was used for final testing. In total, 96 embryos were used: 55 for training, 20 for validation, and 21 for testing of the model. The Adam optimizer was used for loss optimization. Test metrics on the unseen hold-out dataset are shown in Fig. S1d with PSNR 36.94, MS-SSIM 0.94, and PCC 0.81, respectively.

The model framework was based on a library^56^ in Tensorflow 2.3. We trained the model on NVIDIA GeForce RTX 2080 Ti. The training for 10 epochs took 4 hours and 56 minutes.

### Health grading data annotation

Embryos were cultured in culture medium only (untreated), or culture medium containing pure phthalate mixture (treated) to halt growth and mimic a population of poor quality/developmentally arrested embryos. After imaging, z-composite LS-GLIM images (maximum phase projection) of the z-stack were computed for each embryo in the whole sample set to pass on to the embryologists for annotation. One of the three grades: healthy, intermediate, or sick, was assigned to each embryo by one or more expert embryologists. Untreated embryos could therefore fall into any of the three categories. However, treated embryos would be graded as intermediate or sick depending on the severity of damage induced by the treatment. The intermediate label was subjective in that the embryo’s fate was unknown at the time of fixation. The data collected over several rounds of sample preparation and imaging experiments were combined into one collective dataset with a total of 152 embryos graded as 46 healthy, 65 intermediate, and 41 sick. The training, validation, and initial test data were then extracted from this collective dataset individually for each health grading model (see below). An additional test dataset of 43 embryos was subsequently merged with the initial test dataset giving a total of 195 embryos used for model development.

### 3D segmentation and feature extraction

3D nuclei predictions were stacked up into a volume, and the 3D nuclei features were extracted using a 2-step segmentation procedure. The first step of the segmentation involved 2D segmentation per z-section image. Each z-section image of size 1280 x 1280 pixels was median filtered with a window size of 19 pixels in both dimensions to make the intensity uniform and then normalized between its minimum and maximum value. The images were then hard thresholded using a threshold of 0.2, removing any stray detections in the non-nuclei area. Following the hard-thresholding, adaptive thresholding with a sensitivity of 0.55 and a window size of 145 pixels in each dimension was performed to generate a nuclear binary mask. Morphological operations such as opening and area filtering with a cut-off of 400 pixels^2^ (corresponding to a radius of ∼ 1µm) were performed to correct for non-specific detections after binarization. Next, a watershed algorithm was applied to separate nuclei whose borders were touching. After the watershed, other rounds of area filtering were applied to remove elements below the size cut-off of 60 pixels in diameter (∼5 µm), with solidity below 0.8, to remove over-segmented artifacts. The cut-offs of area and solidity were derived empirically after observing the minimum size of nuclei in the images and the associated solidity of the binary mask over the nuclei. Each nucleus in a z-section image of an embryo was assigned a unique label. The exact process was repeated for all z-section images of the entire embryo. The next task was to connect these 2D labels to form 3D labels. Centroids of each nucleus were tracked in the z-direction, and nuclei in two adjacent z-section images were assigned to the same trajectory if their Euclidean distance between the centroids was within a specified sensitivity (50 pixels radius in xy plane). This cut-off was chosen because the two smallest nuclei adjacent to one another would have at least 60 pixels of the distance between their centroids. After assigning centroids to trajectories, we checked for breaks in the trajectory for the cases of nuclei stacked on one another along the z-dimension. To find such separator boundaries, we looked for a series of local minima in the z-direction for the area of the 2D slice of the nuclei and the distance from the first centroid in the trajectory. Gaps in trajectories of more than 5 µm were used as the definition of a new nuclear boundary. Each nucleus in a 3D trajectory was assigned one unique label and was added to the existing embryo 3D label volume. Finally, filtering based on empirically determined minimum volume in voxels (5000 px^3^), maximum volume (250000 px^3^), minimum z-depth (3), and minimum extent (0.14) of the 3D nuclear volume was performed to remove over/under-segmented particles. After 3D segmentation, each nucleus in an embryo volume was assigned a unique color-coded identification label that corresponds to the identity of individual nucleus (Fig.2 f). It is important to note that this 3D segmentation remains in the pixels domain, and rendering at this point will not accurately represent the 3D volume. For proper 3D measurements, the labeled volume was resized by the lateral pixel ratios (11 pixels per µm) and z-step size (1µm), and the labels are interpolated along the z-direction using ‘nearest’ interpolation such that an accurate 3D representation is achieved, while maintaining the labels across the interpolated volume. The phase volume was interpolated with linear interpolation in the z-axis.

The max value projection of the labeled image can then be used for producing a nuclear count map, as shown in Fig. S4, with the color bar denoting the number of nuclei in a single embryo. Features representing nuclear shape descriptors (volume, surface area, sphericity) and nuclear composition descriptors (dry mass and dry mass density) were then extracted for each nucleus. Embryo-level features (nucleus count, scattering amplitude spectrum bandwidth) based on the aggregate system of nuclei inside an embryo were also extracted after 3D segmentation. For subsequent data analysis, nuclei with sphericity greater than 1 were excluded as those represented under-segmented clusters.

We also prepared a normalized dry mass density map per embryo (Fig. 4) based on the extracted nuclear dry mass density. The color of each nucleus represents the mean nuclear dry mass density averaged over its volume. The volume was Gaussian filtered with standard deviations of (1,1,3) for x, y, and z dimensions respectively to enhance the smoothness of the 3D representation. Each embryo was then normalized over the volume for meaningful comparisons between different embryos.

### Scattering amplitude modeling

The distribution of nuclei inside an embryo can be modeled as a system composed of multiple identical repeating units. The scattering potential of such a system can be expressed as^24^

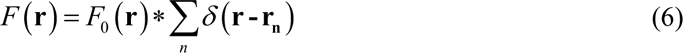

The contribution of individual repeating units is represented by the delta functions placed at different position vectors **r_n_**, * denotes 3D convolution in the spatial domain and *F*_0_ (**r**) is the scattering potential of a single unit.

The scattering amplitude^24^ can then be expressed as the Fourier transform of Eq. (6)

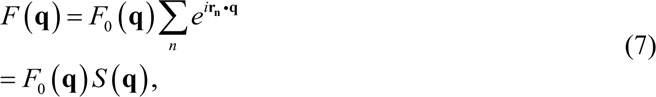

where, **q** is the scattering wave vector.

The two terms of this equation convey different information. The first term *F*_0_ (**q**)is the *form function*, which represents the scattering amplitude’s envelope, while the fluctuations within the envelope are determined by the second term 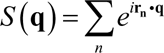, and defined as the *structure-function*. The form function is the Fourier transform of the scattering potential of a single repeating unit and is dependent on the shape of that unit. The structure function is the aggregate effect of all repeating units and is only affected by the distribution of such units inside the system’s volume. For an embryo the repeating units are represented by the individual nuclei. To make the assumption of identical repeating units valid, nuclei were replaced with a unit sphere centered at the centroid of each nucleus. In doing so, the size/shape variability between the nuclei was eliminated. Since we used nuclei predictions to construct the 3D embryo space, we do not have any other part of the embryo, like cell cytoplasm, etc., in our system. The resultant modeled embryo systems for a healthy/intermediate and a sick embryo are shown in

Figs. S2a and S2b respectively. The modeled system’s 3D Fourier transform (a product of form and structure functions) can then determine differences in the embryos based on their health class. A maximum intensity projection along the **k_z_** direction of the scattering amplitude spectrums for the corresponding healthy/intermediate and sick embryos are different (Figs. S2c and S2d respectively), indicating a difference in power distribution. It is important to note that the maximum projection is only shown for visualization purposes, and all the calculations are performed in the 3D spatial frequency space. The raw radial average of the power spectral density of scattering amplitude is shown in Fig. S2e for all 152 embryos, where the red curves are associated with sick embryos, the blue curve for intermediate embryos, and the green curve for healthy embryos. It is immediately observed that the curves for healthy and intermediate embryos overlap. To illustrate the difference between the three classes, the curves were averaged per group (Fig. S2f) and divided by their respective peak powers to get the final normalized radially averaged powers spectral density of the scattering amplitudes for the three classes (Fig. S2g).

To calculate the bandwidths of the scattering amplitude spectrum per embryo, images were downsized in the xy plane to prepare a 3D volume and enable a 256-point FFT in each spatial direction. The frequency space was rescaled by the same factor to get the correct spatial frequencies. After placing unit radii spheres at the centroids of individual nuclei, we performed a 3D FFT to calculate the spectrum. The spatial frequency 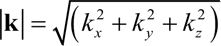 was discretized into 256 levels, and a radial average of the spectrum was calculated on a thin spherical shell covered between successive elements of the **k** vector.

The 3dB or 50% power bandwidth was extracted from the radial average, representing the bandwidth at which the power drops to 50% of its peak power and was used as an essential feature in health grading.

### Health grading of embryos

Based on our analysis of parameters between the healthy, intermediate, and sick embryos, we found that there are insignificant differences between the parameters of healthy and intermediate embryos, while highly significant differences between either healthy and intermediate versus sick embryos (Supplementary Table 3). This information was used to group the embryos into two classes: a combined H/I class (healthy and intermediate embryos) and S class (sick embryos). Our task was to classify each embryos’ health into these two classes-H/I or S. To test the accuracy of health classification based on the extracted features only, we trained a neural network classifier. We called this model a feature-based model (FBM) for health grading. In addition, to take advantage of the state-of-the-art models, we trained a second deep learning model that would not require any hand-engineered features as it will predict the health of the embryos from the LS-GLIM images alone. We called this second model an image-based model (IBM) for embryo health grading.

### FBM architecture and training details

The features used for training were those showing higher statistical significance from the feature pairs with a correlation above 0.6 (for example dry mass-dry mass density and volume-surface area) on the heatmap in Fig. ED1h. Nucleus count (nuc_count) displayed a high negative correlation (∼ −0.7) with scattering amplitude spectral bandwidth (bw3dB), however it was included in the feature vector because it represents the absolute number of nuclei per embryo (Fig. S4). The selected features were arranged in a tabular dataset with five predictors: bw3dB, nucleus count, nuclear dry mass density, nuclear surface area, nuclear sphericity, and two responses 0 (for H/I) or 1 (for S). Data from the 152 embryos were divided such that 102 embryos were used for training, 16 for validation, 34 were kept as holdout data for testing to be combined with the second experimental dataset of 43 embryos for full testing. The initial model type search was performed in MATLAB Classification Learner app with the model type selection set to all models. The models were tested on the combined test dataset as well as on the live test dataset (19 instances of live embryos). For the final common test dataset comparing both the models (FBM and IBM), 5 embryos from the combined test set were not included because they had been used in the validation set of the IBM. The combined test and live performance of neural network classifiers exceed other classifier models (Supplementary Table 4). A neural network-based classifier architecture was therefore selected and optimized to improve the results further.

The FBM is a feedforward, fully connected 3-layer neural network with hidden layers of size [10, 2], constructed and trained using *fitcnet* in MATLAB, with the architecture as shown in Fig. S3a. The hidden layer activations were ReLU and the output activation was SoftMax. Model weights were initialized with Glorot initialization with the biases initialized to zeros. The early stopping is controlled by validation patience, gradient tolerance, and loss tolerance which are set to 20, 1e-9, and 1e-9, respectively. The loss function is the standard cross-entropy loss with L2 regularization. To control overfitting, the regularization term lambda was set to 1e-6.

The limited memory Broyden-Fletcher-Goldfarb-Shanno quasi-Newton algorithm (LBFGS) was employed to minimize the loss function. Although the model’s performance on the hold-out test dataset varied slightly with different random seed initializations, the model performance on the out-of-distribution live dataset varied greatly. The training was rerun using the same model architecture and data, but with a varied random seed (while keeping the random number generator to default: ‘twister’). The model that performed best on the live test dataset with a random seed value of 94202 was ultimately selected.

The loss curve for the model is shown in Fig. S3b, where the training is terminated at the 66^th^ epoch. The final selected model (at epoch=45) has minimum validation loss indicated by the green vertical line.

To determine feature importance, we conducted the Shapley additive explanations (SHAP) test in Python 3.9.7 using SHAP API. The MATLAB model parameters were exported to Python to use the SHAP Python API.

### IBM architecture and training details

The IBM architecture is shown in Fig. S3c. It was specified as a standard Efficient Net B7 architecture with pretrained weights downloaded from the Pytorch-torchvision models package. The classification head was replaced with a 3-layer classifier of shape [2560, 500, 200] with two classes as output and the dropout rate in the classifier was changed from the default 0.5 to 0.3. The model requires the inputs to be in shape [633, 600]; hence, images were downscaled from 1280 x 1280 to 633 x 600. Random horizontal flips (p=0.8), random vertical flips (p=0.7), and random rotations from −30 to +30 degrees were used to augment the training data. Augmentation was performed using the torchvision transforms. The input images to the model were three-channel images called z-slices, with each channel being a neighboring z-section to the central image. The 3-channel image was normalized between the minimum and maximum value of the entire 3-channel stack.

All the model layers except the last convolutional layer (layer 8.0) and the subsequent classifier head were fixed for training. The model was trained using the standard cross-entropy loss with a learning rate of 5e-6. In addition, we employed exponential learning rate decay with a gamma value of 0.9. The standard Adam optimizer with all default parameters except the learning rate was used for loss optimization. The training batch size was set to 4 with a validation batch size of 1. The model was trained on 3809 images (52 embryos) and validated on 1297 images (21 embryos). The remaining images (79 embryos) were kept as a holdout for testing to be combined with data from other subsequent experiments (43 embryos) to create a test set randomized over the multiple cycles of experiments. The loss curve for the model training is shown in Fig. S3d. The loss converged, and the model training was stopped after 100 epochs after no significant improvements in the loss were observed.

We employed Grad-CAM^45, 46^ to detect decisive features in the image. The Grad-CAM maps were extracted after the last convolutional layer of the Efficient-Net B7. The model and associated codes were developed in Python 3.9.7 and Pytorch 1.11.0 and trained on NVIDIA GeForce RTX 3060 Ti. The training for 100 epochs took approximately 48 hours.

### Max-voting procedure

For both the FBM and the IBM, max-voting of individual nuclei/z-slice image predictions was performed to get the overall embryo-level predictions, meaning that the class which was predicted the most for the overall embryo was selected as the final class. Confidence probability was then the average of prediction scores over the majority class subset of nuclei/z-slice image for FBM, IBM respectively.

### Automation of workflow using the MATLAB app

All the MATLAB-based processing codes were combined to develop a standalone MATLAB app to demonstrate our tool. As the MATLAB license terms dictates, this app is only for scientific demonstration with no commercial/monetary usage. The app has three panels. The first is to decide the segmentation parameters for initial 2D and final 3D segmentation based on nuclei predictions, as shown in Fig. S6. After segmentation, the second panel (Fig. S7) analyzes features and performs health grading. The third panel (Fig. S8) is for demonstrating sparse predictions on a random set of z-slices. The detailed protocol is discussed in the supplementary note2: MATLAB standalone app operation.

### ROC fitting and standard error estimation

The ROC curves in Fig. ED2 were fitted to extract standard error of the mean (sem) values using the ROC analysis tool developed by Metz et al. ^57^. We chose semi-parametric estimation with a conventional binormal ROC curve model. The inverse of the information matrix was selected as the uncertainty estimation method.

### Statistical analysis

All the extracted features were first tested for normality using Lilliefors Test (at 5% significance). All the features except nucleus count and bw3dB were found to be non-normal (Supplementary Table 3). Kruskal Wallis non-parametric test (at 0.1% significance) was applied to test for differences in the distribution between the three classes: healthy (H), intermediate (I), and sick (S). Post-hoc Dunn-multiple comparison test with Holm p-value adjustment was applied to determine the pairwise differences in the distribution (Supplementary Table 3). The nucleus count and bw3dB were normally distributed thus the Levene test was applied to test for equality of variance. The variances were similar (p>0.001, the null hypothesis was not rejected). We then applied one-way ANOVA and Student T-test post-hoc test with Holm adjustment to test for deviations in group means. Results are shown in Supplementary Table 3.

All the statistical analysis was performed in Python 3.9.7. In addition, we used the following libraries for individual tests: Lilliefors (statsmodel api), Kruskal-Wallis test, Levene test, one-way ANOVA test (scipy stats), Dunn’s posthoc, post-hoc Student T test (scikit posthoc).

### 3D renderings

Amira (Thermo Fisher Scientific) was used for the renderings shown in Figs. 2 d, e, and supplementary movie 2. Clear volume (Fiji, ImageJ, NIH) was used for Fig. 2f. MATLAB 2022b (MathWorks) was used for all other 3D renderings.

## Data availability

All the data generated in this study are included in the manuscript. Imaging data are not shared due to size constraints.

## Code availability

Standalone MATLAB app, Python scripts for inference and the three models used/developed in this study will be uploaded to Github https://github.com/NehaRG-QPI/embryo_ls_glim.

## Supporting information

Supplementary Movie M1: z-scan of an embryo with overlay of LS-GLIM (gray) and nucleus predictions by NPM (red).

Supplementary Movie M2. 3d visualization of stacked ground truth (fluorescence) and stacked nuclei predictions on a test embryo for comparison.

Supplementary Movie M3. Movie depicting 3D renderings of LS-GLIM, nuclei prediction, segmentation labels, and mean nuclear dry mass density map

## Acknowledgements

The authors of this study acknowledge Yuchen R. He for the development of Efficient-Net encoded UNet model used in this study for nucleus detection. We also dedicate this work to Prof. Gabriel Popescu, who tragically passed away during the advanced stages of the study. His passion for science and deep insights in phase imaging will be dearly missed.

## Author contributions

G. P., R. A. N., and N. W. conceptualized the project. N. G. designed the imaging experiments. N. Z. E. L., R. B. A., and W. C. cultured, stained, and prepared the embryo samples. X. C. selected nucleus detection stain 7-AAD. R. A. N., and N. W., graded the embryos for preparing ground truth data. N. G. conducted imaging experiments, data analysis, trained machine-learning models and did the post-prediction analysis. N. W., R. A. N., and W. C. were the experts in the blind analysis for image-model interpretation. N. S. supervised initial IBM model development. R. A. N., N. W., G. P., and M. A. A. supervised the study. N. G., R. A. N., N. W., and M. A. A. wrote the manuscript with contributions from all authors.

---

**Extended Data Figure 1.**
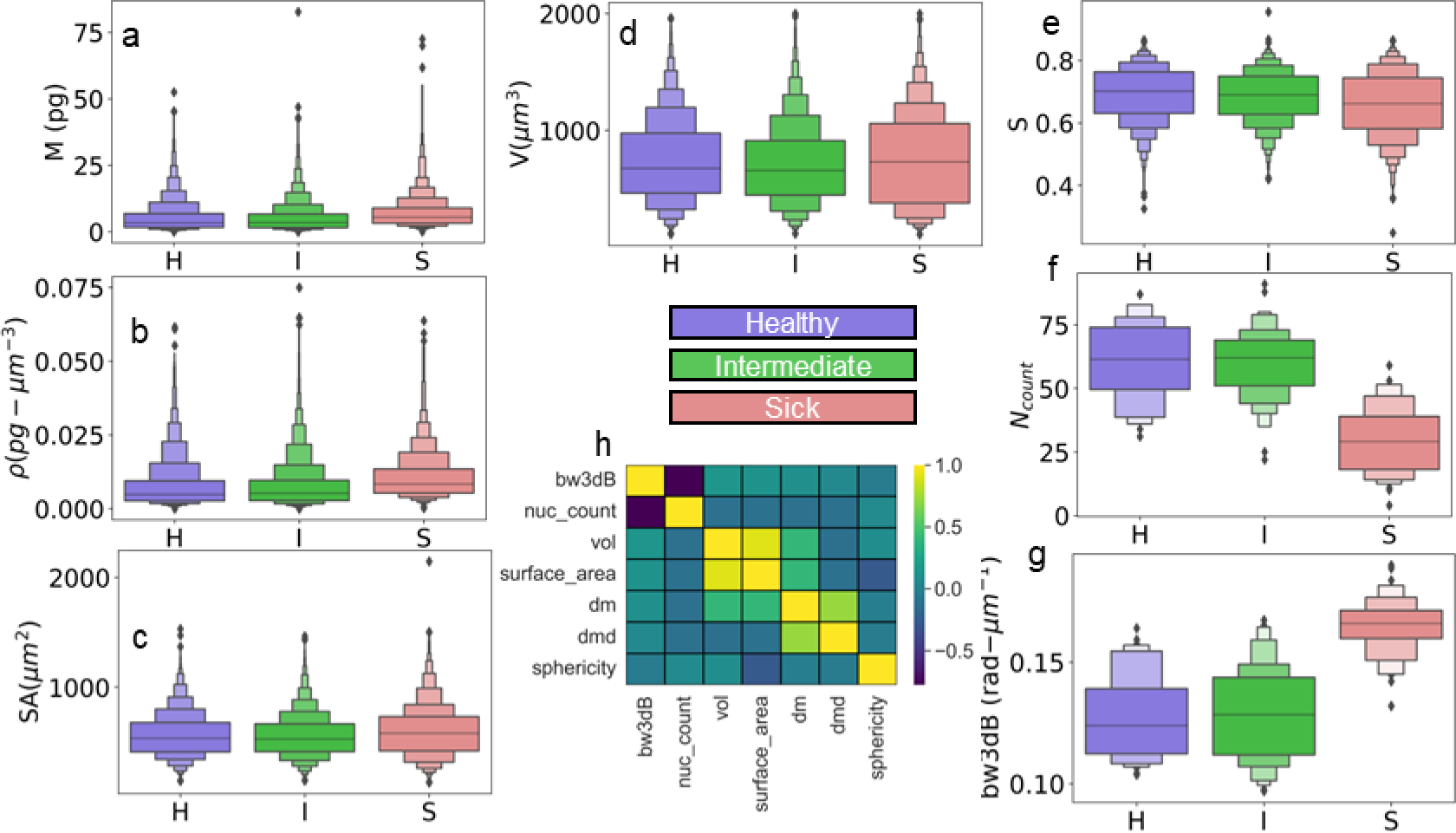
Features: Boxenplots for a. Dry mass b. Dry mass density c. Surface area d. Volume e. Sphericity f. Nucleus count g. bw3dB for healthy (blue), intermediate (green) and sick (red) embryos. h. Correlation heat map for all features.

**Extended Data Figure 2.**
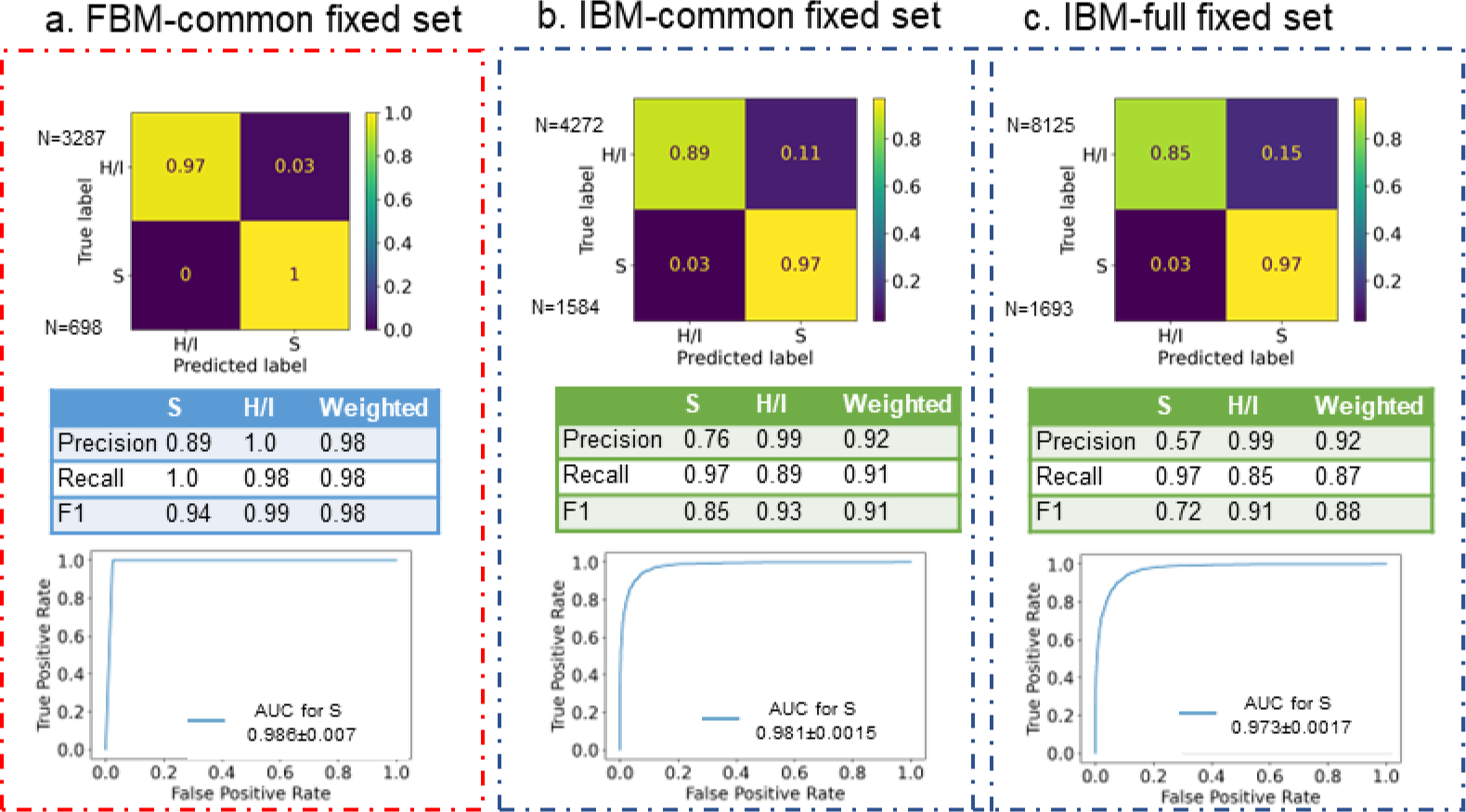
Health grading baseline model performance-confusion matrix, performance metrics, ROC curve for in-distribution test set. a FBM (nuclei level-72 embryos) b. IBM (z-slice level-72 embryos), c. IBM (z-slice level-122 embryos).

**Extended Data Figure 3.**
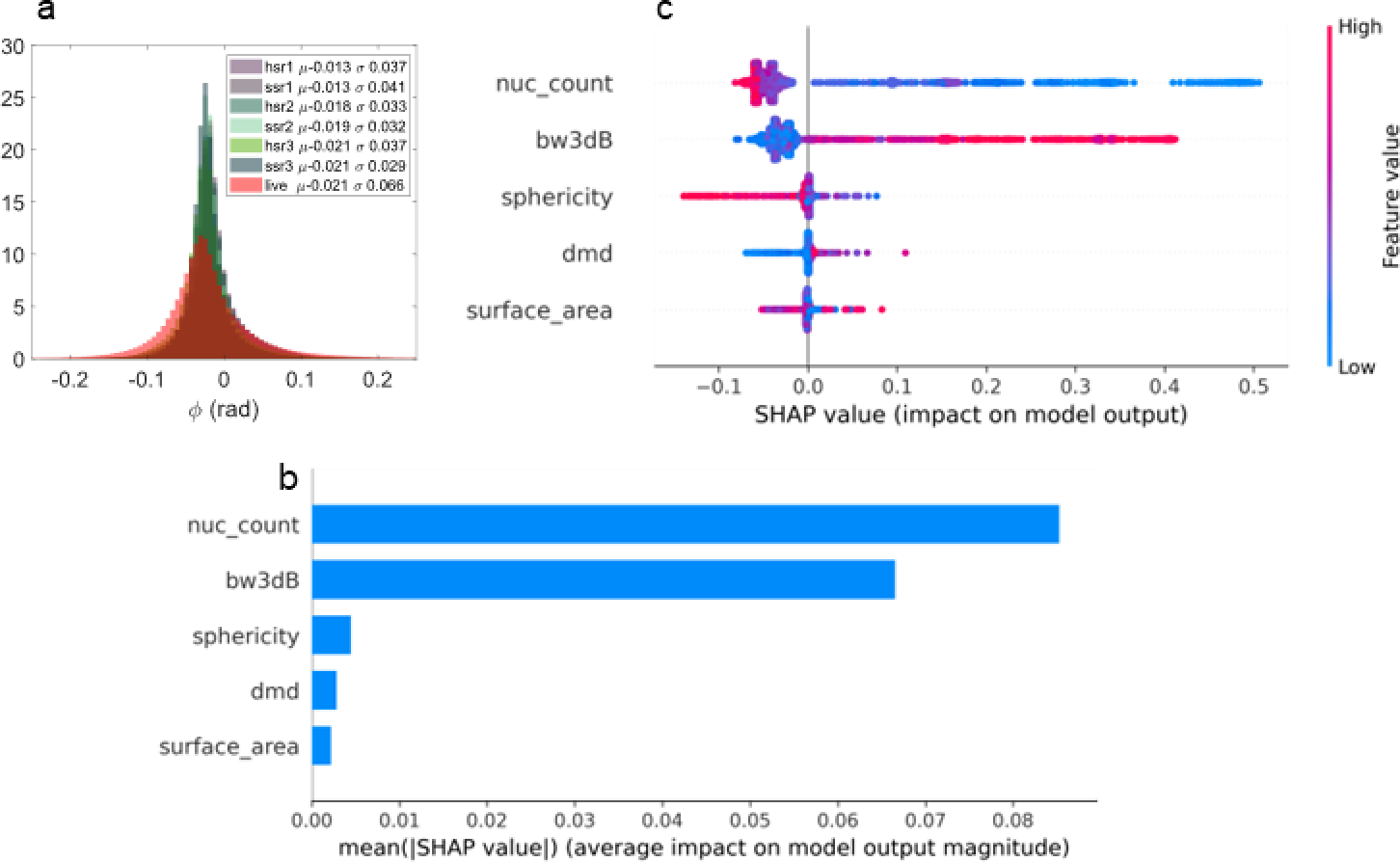
a. Phase distribution histogram between in-distribution datasets (hsr1, ss1, hsr2, ssr2, hsr3 and ssr3) versus out-of-distribution dataset (live). b. SHAP feature importance plot for FBM, c. SHAP summary plot for FBM

**Extended Data Figure 4.**
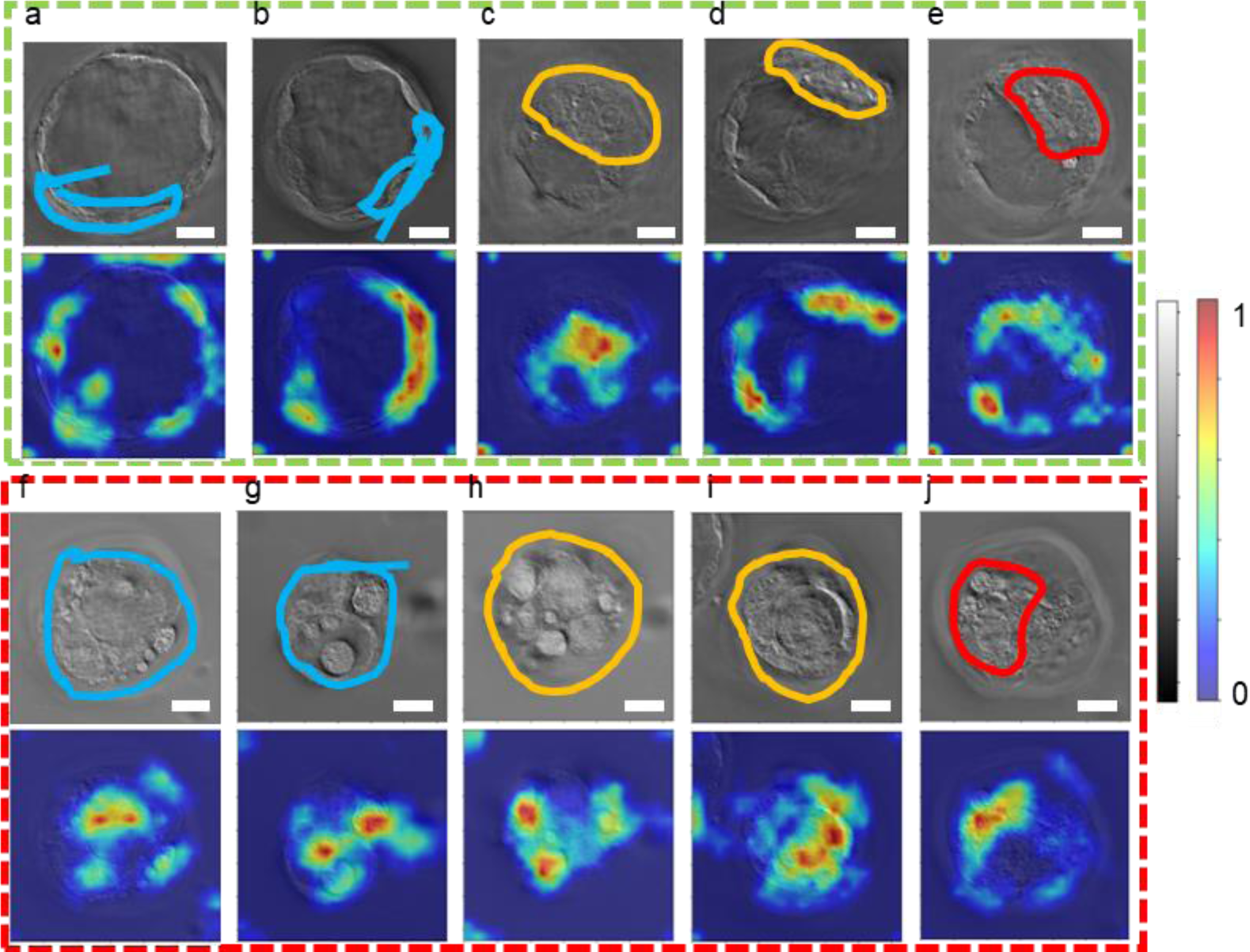
Grad-CAM interpretability test of IBM: for healthy (a-e) and sick (f-j) embryos for successful predictions: The top row in each panel is a random z-slice of different embryos with the corresponding overlayed Grad-CAM image in the bottom row. Results are compared with blind markings from three experts (shown with different colors). Colormap for GLIM images shows normalized phase maps and the color bar for Grad-CAM outputs shows the importance of the regions for correct prediction with red being the highest.

**Extended Data Figure 5.**
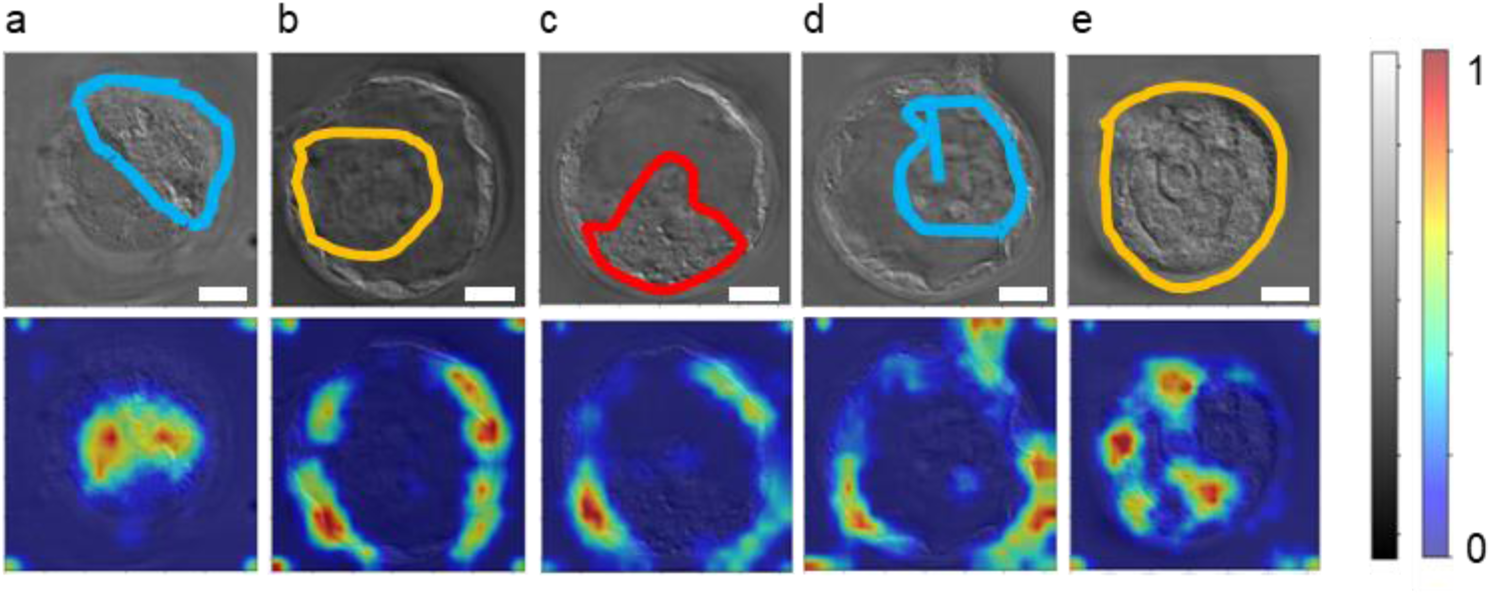
Grad-CAM interpretability test of IBM: for healthy (a-e) embryos for successful predictions where expert decisions do not fully agree: The top row in each panel is a random z-slice of different embryos with the corresponding overlayed Grad-CAM image in the bottom row. Results are compared with blind markings from three experts (shown with different colors). Colormap for GLIM images show normalized phase maps and the color bar for Grad-CAM outputs shows the importance of the regions for correct prediction with red being the highest.

**Extended Data Figure 6.**
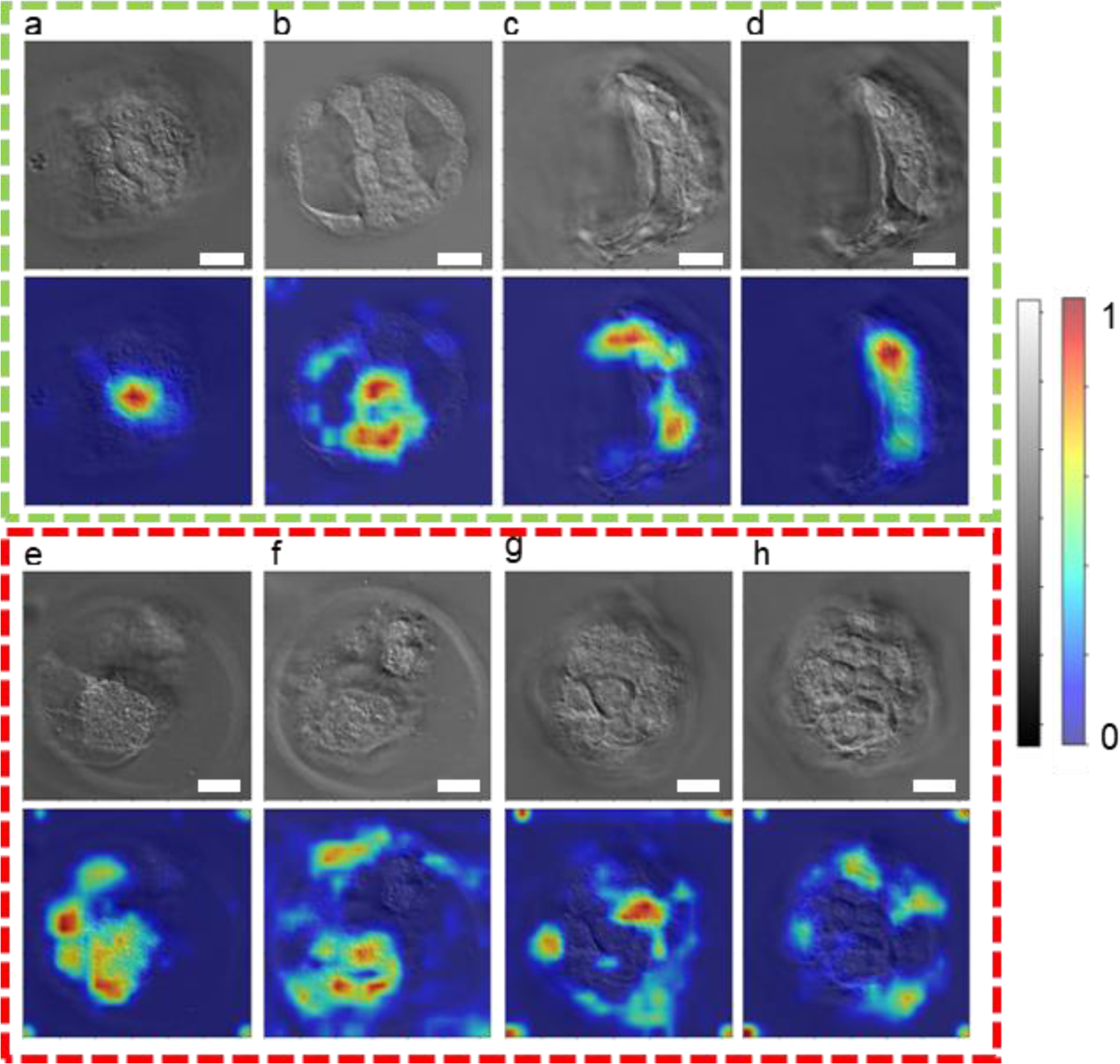
Grad-CAM interpretability test of IBM: for wrong predictions: Healthy z-slices predicted as sick (a-d), sick z-slices predicted as healthy (e-h). Pair of slices per embryo are shown such that a and b belong to one embryo, c and d belong to another embryo, e and f belong to another embryo, and g and h belong to another embryo.

**Extended Data Figure 7.**
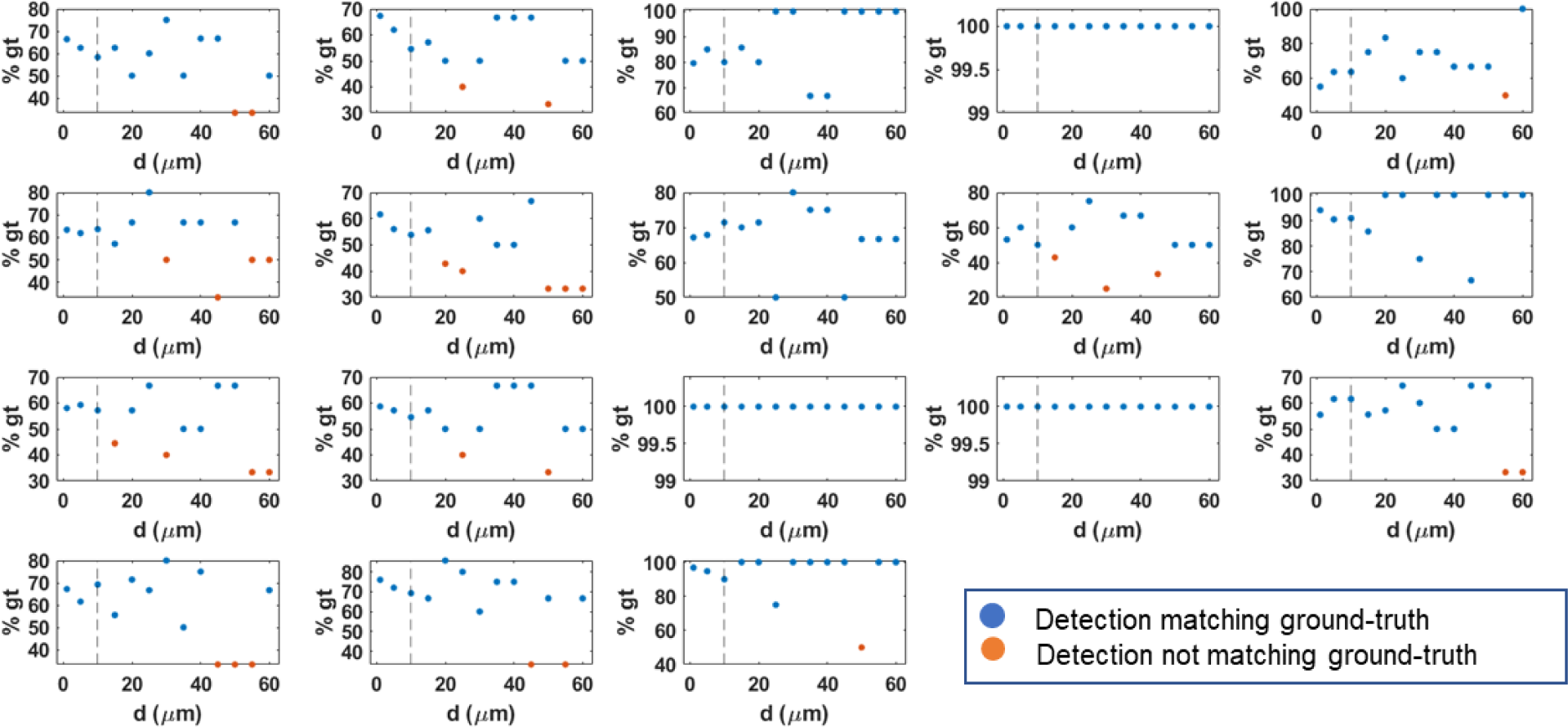
IBM evaluation of sparse predictions for maximum z-slice interval: Each plot represents one live embryo instance. z-slices for each embryo were selected according to the increasing interval between them starting from 1µm to 60 µm (x-axis of each plot). Correspondingly, the % of the detected class is plotted on the y-axis. The black vertical line on each graph shows the maximum interval (10 µm) after which error starts to appear. Blue dots are correct predictions (matching to the ground truth) and an orange dot appears when the model flips the prediction to the wrong class due to insufficient z-slices.

**Extended Data Figure 8.**
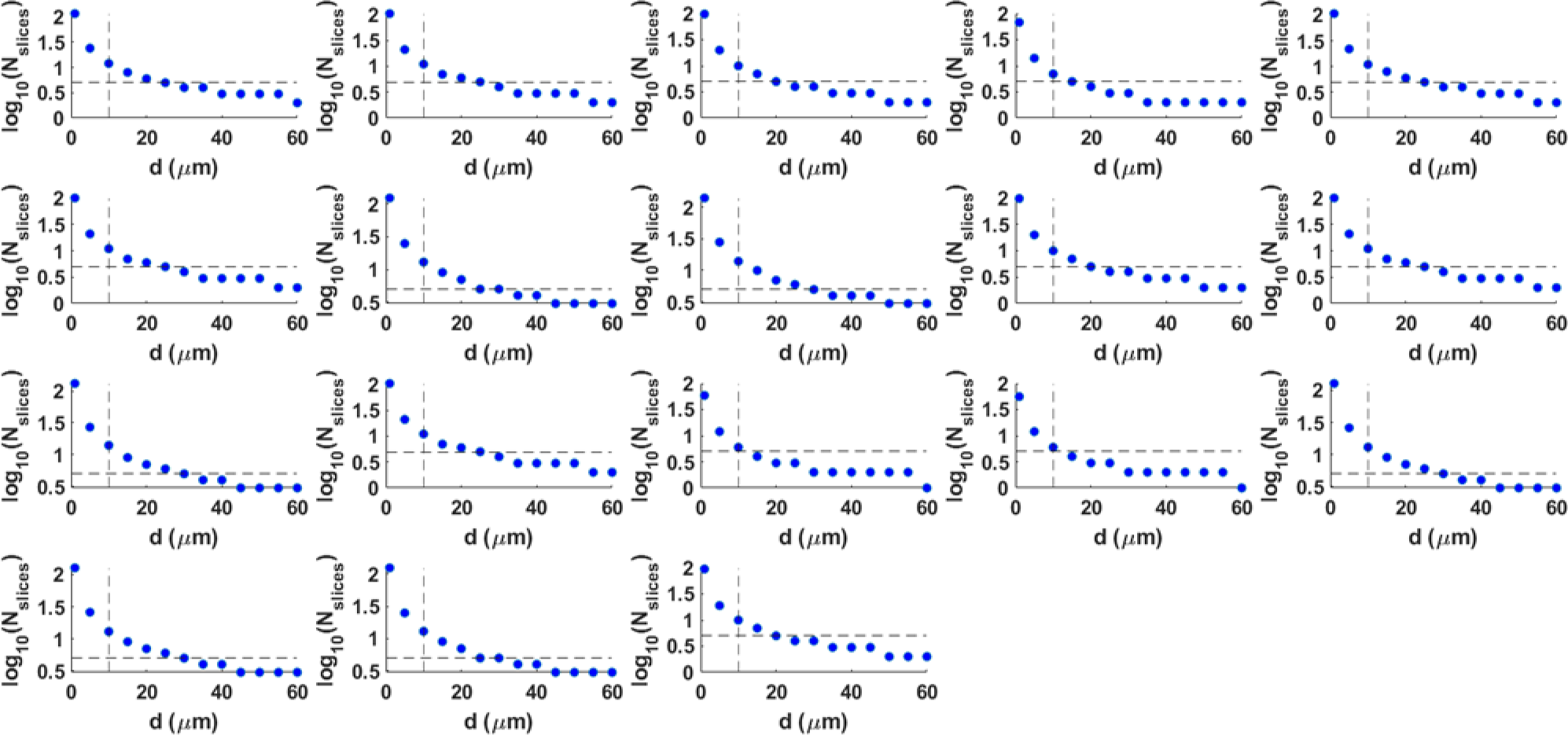
IBM evaluation of sparse predictions for minimum number of z-slices: Each plot represents one live embryo instance. z-slices for each embryo were selected according to the increasing interval between them starting from 1µm to 60 µm (x-axis of each plot). The corresponding number of z-slices to cover the entire embryo are shown on the y-axis in log10 scale. Black vertical line is the maximum interval (10 µm) determined from extended data figure 7. For intervals less than 10 µm, the number of z-slices are greater than 5, shown by the dotted black horizontal line.

**Extended Data Figure 9.**
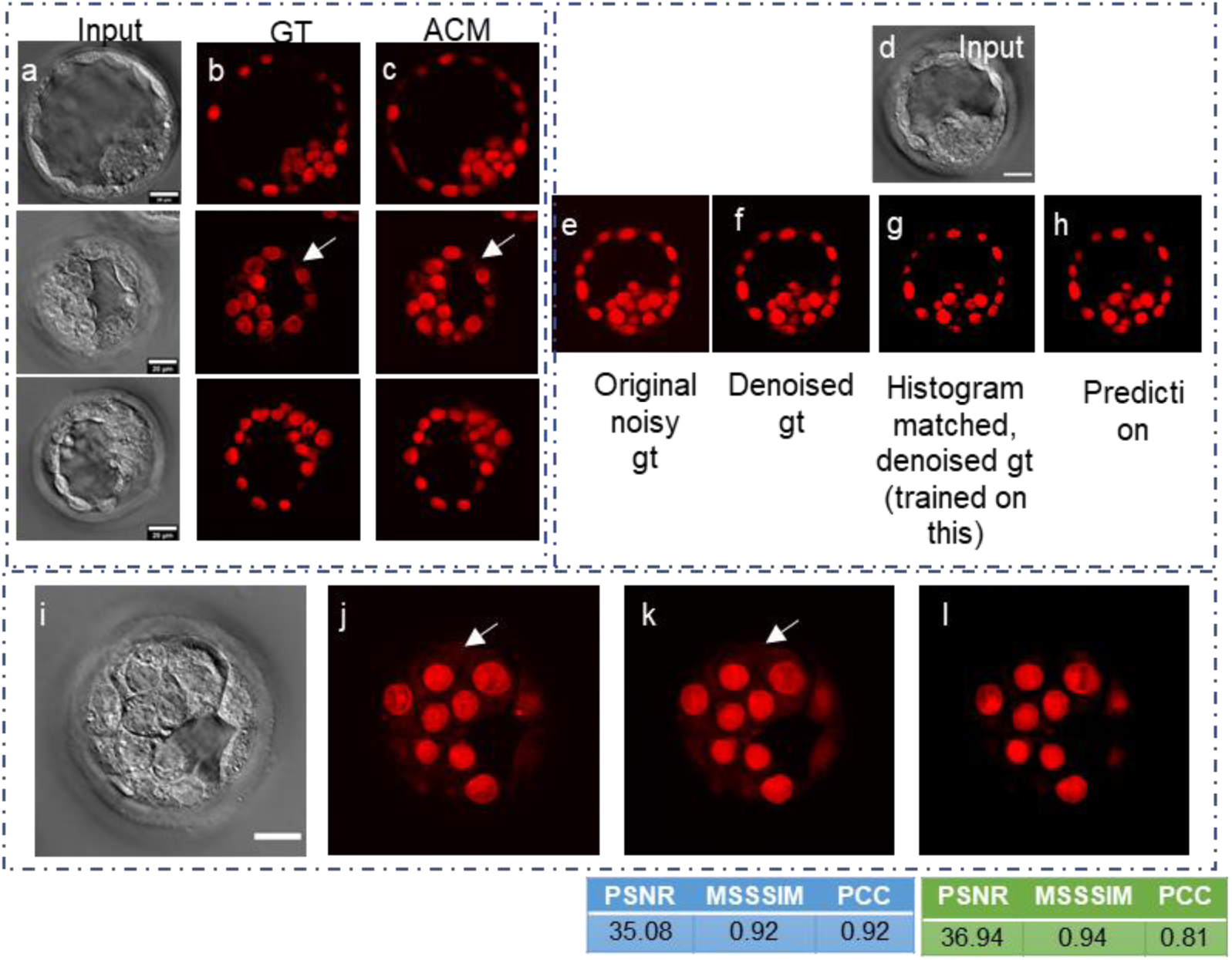
Nucleus prediction model preprocessing. a. Input: LS-GLIM, b. ground truth fluorescence, and c. model predictions without histogram matching. Preprocessing for the final model: d. input LS-GLIM, e. original ground truth, f. denoised ground truth, g. histogram matched to input ground truth, and h. the final prediction by training on images like (d, g) pair. i. one LS-GLIM image, j. Real denoised fluorescence ground truth k. Non histogram-matched Model predictions, l. predictions by final model trained on histogram matched denoised ground truth. White arrows indicate spurious cytoplasm signals. Scalebar is 20 µm for all images.

