## Supplementary text for "Machine learning assisted health viability assay for mouse embryos with artificial confocal microscopy (ACM)"

*Deceased

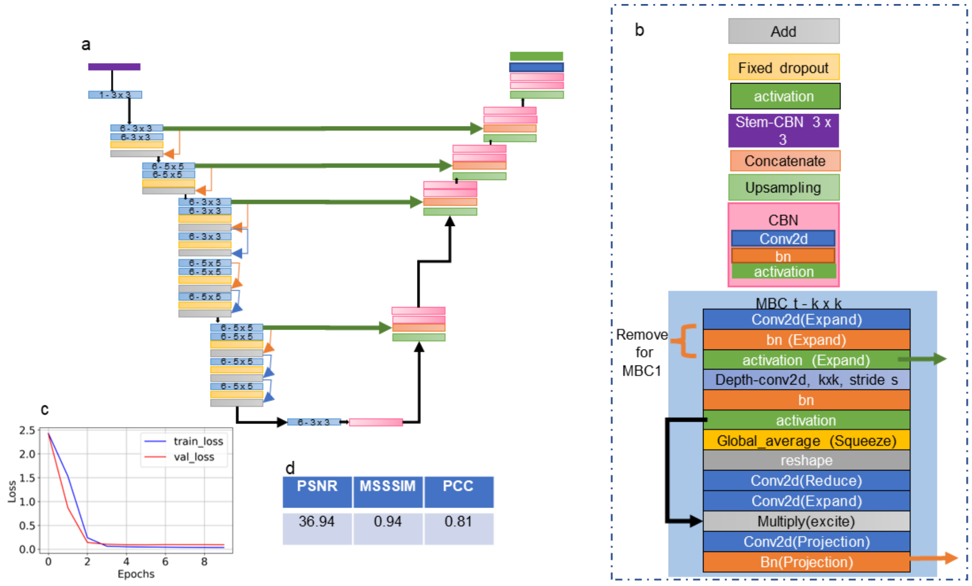


**Supplementary Figure S1. Nucleus detection model architecture** a. Model architecture of Efficient Net-B0 UNet used for nuclei detection. In this model, a pretrained Efficient Net B0 block replaces the encoder part of the UNet. b. Block descriptions of a, c. Loss curve for the model, d. Performance metrics on the test dataset.

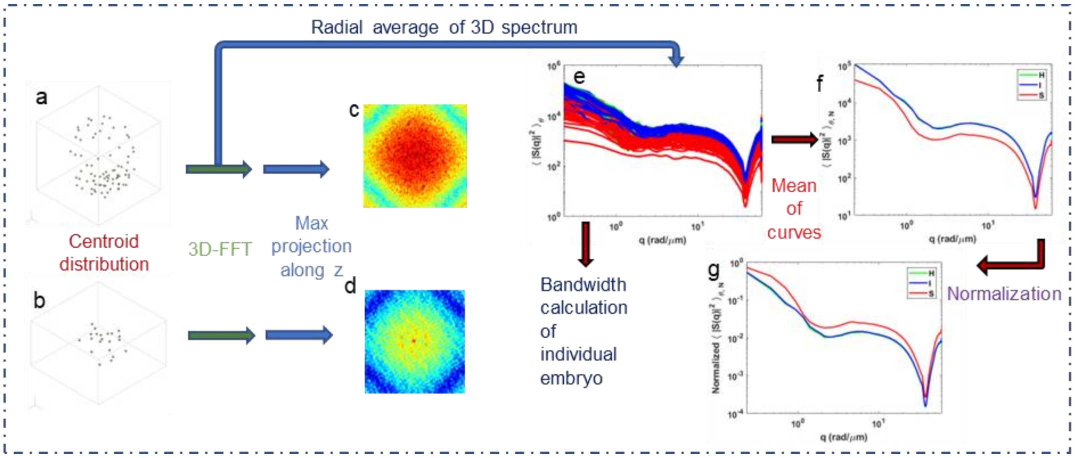


**Supplementary Figure S2. Scattering amplitude bandwidth calculation:** a, b. Centroid distribution from the 3d segmentation, where a unit sphere is placed at each centroid for healthy/intermediate (a) and sick embryo (b). Centroid distribution is then 3D Fourier transformed into $\mathbf{q}$-domain. c and d show the maximum projections of scattering amplitude spectrums along the $\mathbf{k}_z$ direction for visualization purposes only. Note that the calculation of associated bandwidths considers whole 3D volume information and not max projections. e. Radially averaged power spectral density of scattering amplitude associated with all 152 embryos, f. Mean of curves per class, g. Normalized mean radially averaged power spectral density for each class. Red curves denote sick class, green curves denote healthy class and blue curves denote intermediate class.

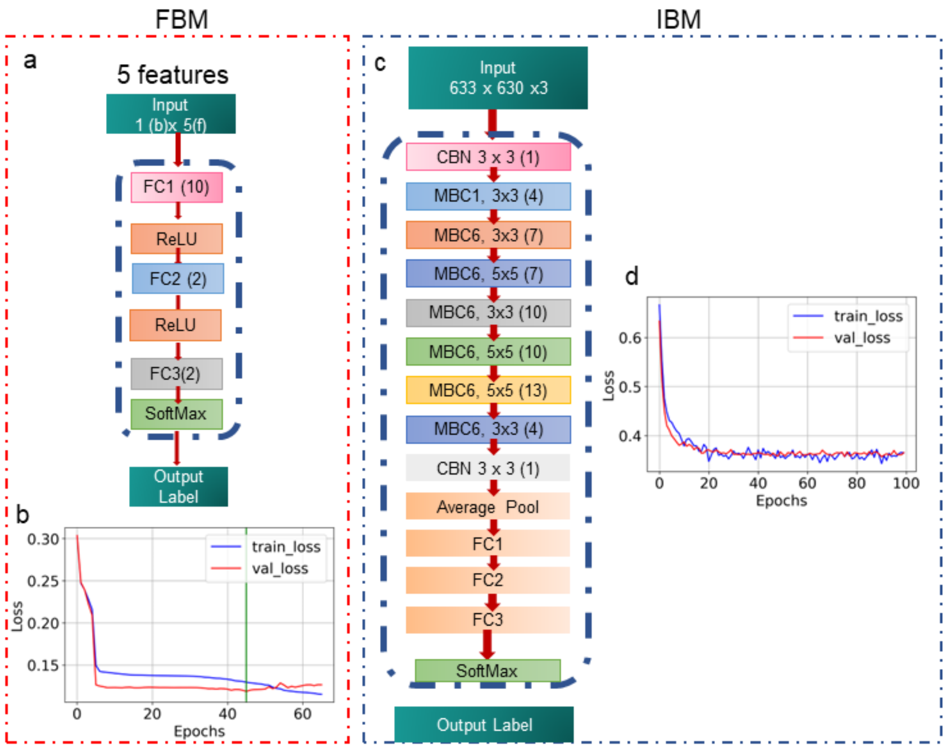


**Supplementary Figure S3. Health grading networks architecture:** a. FBM architecture- a three layered neural network classifier with hidden units [10, 2] b. Loss curve for (a) with a vertical green line denoting the chosen model. c. IBM architecture-an Efficient Net B7. The classifier layer of the model is replaced with three fully connected layers for 2-class classification Corresponding loss curve is shown in d. CBN: Conv2d, batch normalization, activation, MBC: MBConv Block, FC: Fully connected layers.

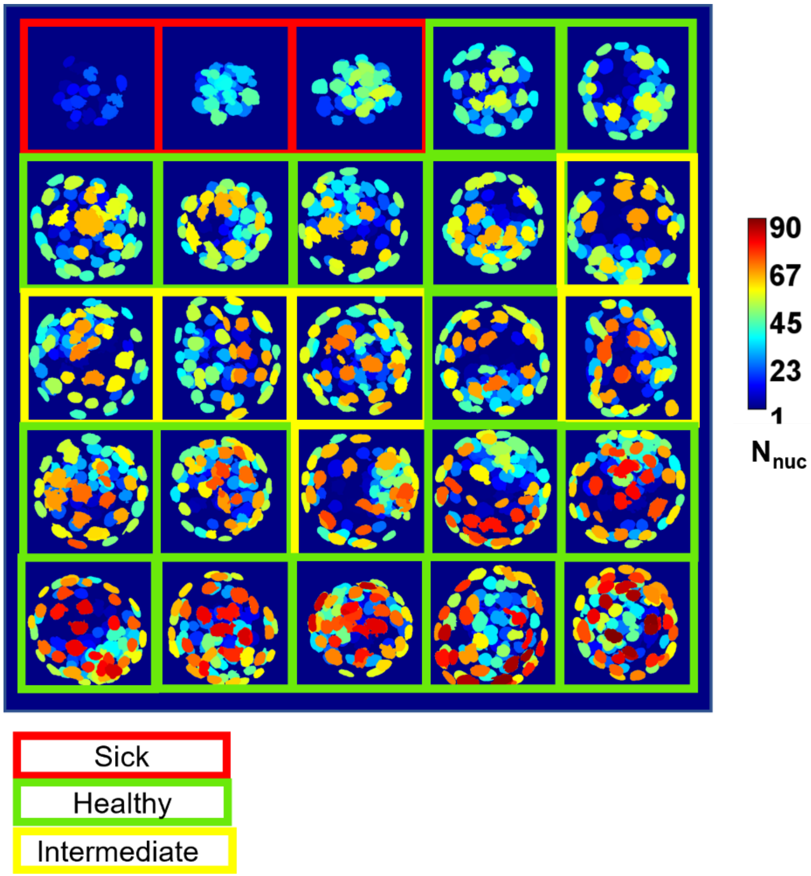


**Supplementary Figure S4.** Nucleus count distribution for a batch of embryos, with embryos in red, green, and yellow boxes belonging to classes Sick, Healthy, and Intermediate, respectively. Colorbar shows the nucleus count distribution.

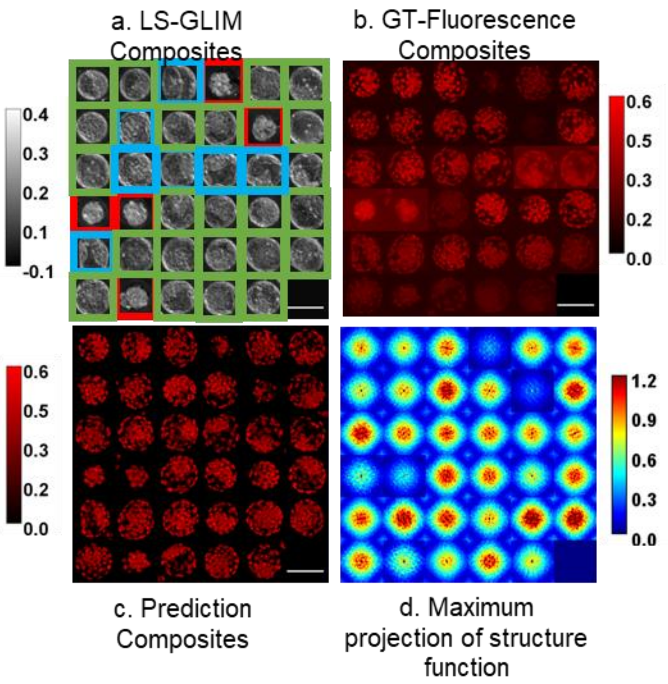


**Supplementary Figure S5. Example of differences in scattering amplitude spectrum for healthy/intermediate and sick classes**. a. LS-GLIM maximum phase projections along z-axis with red boxes showing sick embryos, green boxes showing healthy embryos and blue boxes showing intermediate embryos. Colorbar represents phase distribution b. Ground truth maximum intensity projections of nuclei fluorescence image corresponding to embryos in a, Colorbar represents normalized intensity c. Maximum intensity projections of nuclei model predictions, Colorbar represents normalized intensity d. maximum intensity projections of scattering amplitude spectrum for embryos in a. Colorbar represents spectrum intensity. Scalebar is 100 µm for a, b and c.

**Supplementary Note 1: LS-GLIM setup**

Light from the laser source is directed through the beam steering components inside the LSM box and scanned through the transverse plane of the sample using scanning mirrors SM1 and SM2. The microscope is configured in a differential interference contrast (DIC) mode with a Nomarski prism (NP) beneath the objective that divides the incoming light beam into two laterally sheared orthogonal polarizations. These laterally sheared orthogonal polarization beams travel through the sample and are collected by a condenser lens (CL). After passing through a second Nomarski prism in the condenser, the two laterally sheared beams are combined into a single beam. This composite beam now enters the GLIM module that comprises a liquid crystal variable retarder (LCVR) to introduce modulations in phase. The LCVR provides four phase shifts in steps of $\pi/2$ between the two orthogonal polarization components of the beam. An analyzer (A) placed at 45 deg to either polarization enables interference which is detected by the transmission photomultiplier tube (T-PMT). After recording four intensity images at the T-PMT, a phase-shifting reconstruction algorithm is applied to extract the phase gradient[1,2]. The final phase image, shown in Fig. 1b, is obtained by integrating the phase gradient image along the shear direction[2]. The corresponding fluorescence image (Fig. 1c) for the same field of view is captured through the confocal detector (CD) in the epi-illumination light path after passing through a pinhole (P) and an appropriate emission filter on the emission filter wheel (EFW) as shown in Fig. 1a.

**Supplementary Note 2: MATLAB standalone app operation**

The app is a demonstration of our workflow for segmentation of nucleus prediction images in 2D as well as 3D, visualization and analysis of the nuclei/embryo descriptor features and prediction of the health class of an embryo using the trained FBM or IBM. For testing, we provide datasets for two embryos, one in H/I class and the other in S class. There are three panels in the app: First panel titled 'Segment' is for the segmentation of the nucleus prediction results. The second panel titled 'Analyze' is for feature extraction and health grading using FBM or IBM. The third panel is titled 'Analyze on few slices', this panel is for health grading of live embryos using sparse predictions by IBM.

We will now discuss each of these panels in detail:

**Segment Panel**

The layout of this panel is shown in supplementary figure S6.

This panel is for 2D (Fig. S6a) and 3D (Fig. S6b) segmentation of the nucleus predictions of an embryo.

Required inputs in Fig. S6 are:

1. **pth**: is the system path of the main folder, with the required subfolders to be named as
    a. **glim**: for storing raw GLIM images from LS-GLIM acquisition. This folder is to be generated by the user.

b. **cropped**: for storing 1280 x 1280 sized cropped images from glim folder. This folder is to be generated by the user.

c. **new**: for storing nucleus prediction images of cropped folder images. This folder is to be generated by the user and the nucleus prediction is performed outside the MATLAB environment.

d. **overlapped**: for storing 3-channel z-slice images for IBM predictions. This folder is generated by the app.

e. **2d_3d_segmentation_final**: for storing segmentation data with subfolders: **'label3'** (2D labels of 3D volume), **'seg_3d_3'** (3D labels of 3D volume), **'seg_centroid3'** (centroid distribution of downsized volume) and **'data_resize'** (extracted 3D features).

2. **med_filter_window:** size of the initial 2D median filter, the filter will be applied to each 2D image.
3. **hard_threshold:** First threshold after normalization of 2D image.
4. **sensitivity:** sensitivity of adapthresh MATLAB function.
5. **adapt_wind:** Neighborhood size parameter of adapthresh MATLAB function.
6. **h:** connectivity parameter of imextendedmin MATLAB function, can be 4 or 8.
7. **solidity:** the solidity cut-off of the detected nuclei objects.
8. **image_size:** size of the square input image.
9. **min_debris pixels:** maximum size of discarded objects before the watershed operation in pixels.
10. **min_area pixels:** Objects with area less than the min_area will be discarded after the watershed operation. Objects with area >= 1.5 times min_area and solidity greater than solidity cut-off will be selected for final mask. pixel_ratio: pixels per $\mu m$ for the xy dimension.
11. **objective_NA:** numerical aperture of the objective lens used for imaging.
12. **imaging _z_step:** step size of the z-scan in $\mu m$.
13. **lambda:** wavelength for LS-GLIM acquisition in $\mu m$.
14. **Volume_gating:** Maximum volume cut-off to remove undersegmented nucleus in $\mu m^3$.
15. **gap_z:** Maximum allowed gaps in trajectory in z-direction
16. **nuc_radius:** parameter for determining trajectory of centroid in z-direction.

17. **sensitivity:** sensitivity*nuc_radius determines the amount in pixels in lateral dimensions to search for the next centroid in the adjacent z-sections.
18. **slices to exclude:** use this parameter if the first and last z-sections of the acquired stack do not contain in-focus information. Processing will then be applied to the rest of the z-sections after excluding the specified number of z-sections from top and bottom of the stack.

The test button runs the 2D segmentation on one randomly selected z-section nuclei prediction image and displays the raw image and the corresponding label map in figure panels marked as Fig. S6 c, d respectively. run all button runs the 2D segmentation on whole data and saves the corresponding labels in *pth→2d_3d_segmentation_final→label3* folder.

For the 3D segmentation, test3D runs the 3D segmentation and shows the 3D predicted nuclei volume and the corresponding label volume in Fig. S6 e and f respectively without storing intermediate outputs. This feature is useful for a quicker assessment of 3D segmentation parameters as compared to a full run. Run button performs the 3D segmentation and writes the 3D labels in *pth→2d_3d_segmentation_final→seg_3d_3* and the resized centroid volume in *pth→2d_3d_segmentation_final→ seg_centroid3* folders respectively. The show 3D labels button displays the already stored 3D predicted nuclei volume and the corresponding label volume in Fig. S6 e and f respectively.

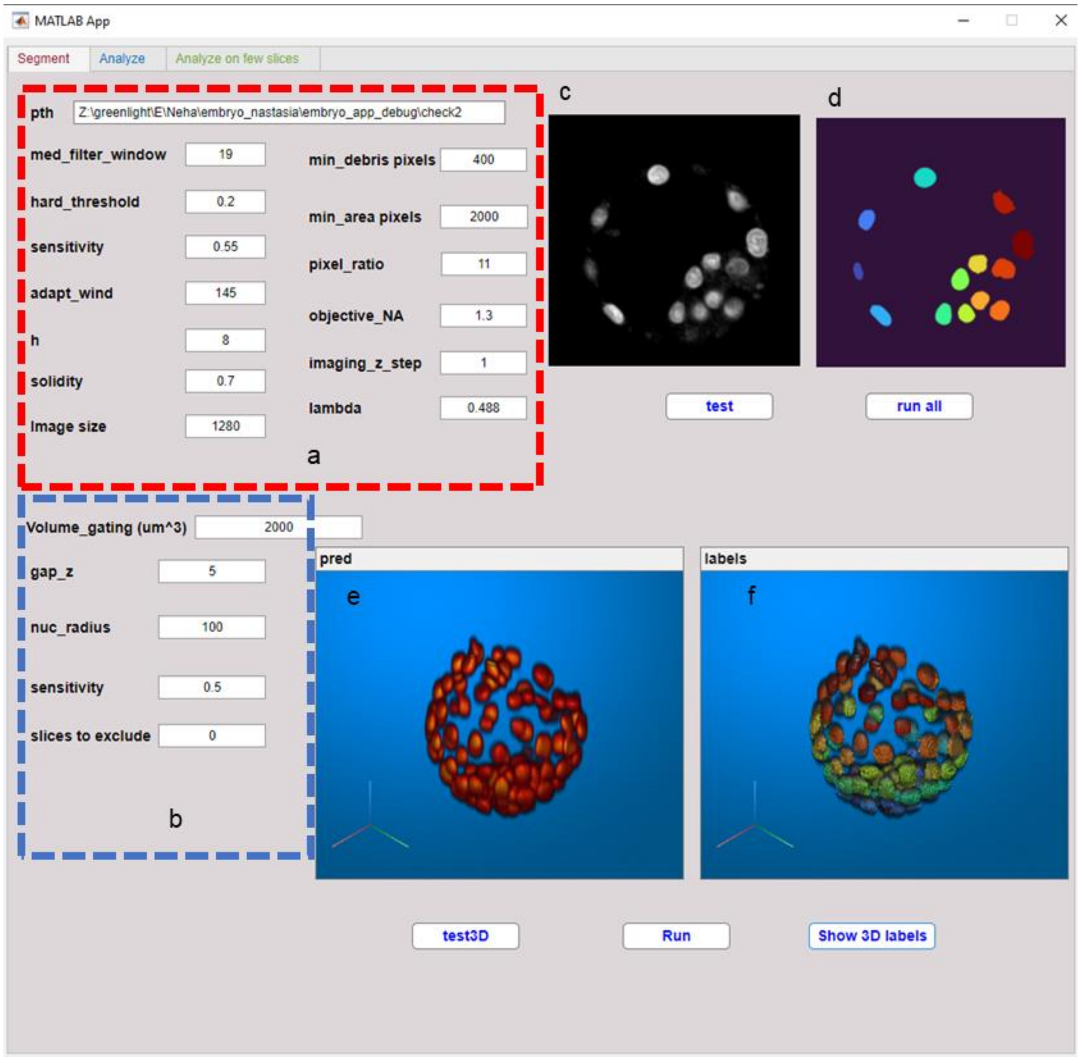


**Supplementary Figure S6. MATLAB standalone app panel 1: Segment**. User defined parameters for a. 2D segmentation b. 3D segmentation c. One z-section of nucleus prediction d. Corresponding generated 2D nuclei instance segmentation label e. 3D stacked nuclei prediction f. 3D nuclei instance segmentation map with each nucleus represented by different colors.

### Analyze Panel

This panel is used to extract 3D features from the segmented nuclei volume of an embryo and perform its health classification. Required inputs in Fig. S7a include:

1. **base_path:** same as **pth** parameter in Fig. S6a.
2. **pth_1d_model:** system path to the FBM (trained in MATLAB).
3. **model_name:** name of the trained FBM model mat file.

The rest of the inputs are the same as in Fig. S6 explained previously. The Exclude slices input is for IBM predictions.

Show composite glim and nucleus prediction button shows the maximum value projection along z-direction of the LS-GLIM and the corresponding nucleus prediction volume (Fig. S7b). Show 3D labels button displays the nuclei prediction volume and the corresponding label volume (Figs. S7 c and d respectively) and also displays the number of nuclei detected in the embryo. Extract parameters and predict-feature model button performs the 3D feature measurements of the labelled volume and classifies the health of the embryo using trained FBM. Show Nuclear dry mass density map button displays the mean nuclear dry mass density distribution map (Fig. S7e) with the colorbar below indicating the values in pg/$\mu m^3$. Based on the FBM output, green LED lights up against H/I-feature text if the embryo is predicted to be of H/I class or a red LED lights up against the S-feature text if the embryo is predicted to be of S class, with the percentage of nuclei in favor of the predicted health class denoted by the slider (Fig. S7f). The histograms of all calculated features are displayed in Fig. S7g. Prepare 3-slice data prepares 3-channel z-slice data from the **cropped** folder and writes the data in the **overlapped** folder. IBM predictions are performed outside MATLAB environment. IBM outputs a **check.csv** file that is saved in the main base folder (**pth**). Predict-Image-model performs the max-voting on individual z-slice predictions of IBM from the check.csv file and the health classification and the percentage of z-slices in the favor of predicted class are displayed in Fig. S7h in a manner similar to Fig. S7f.

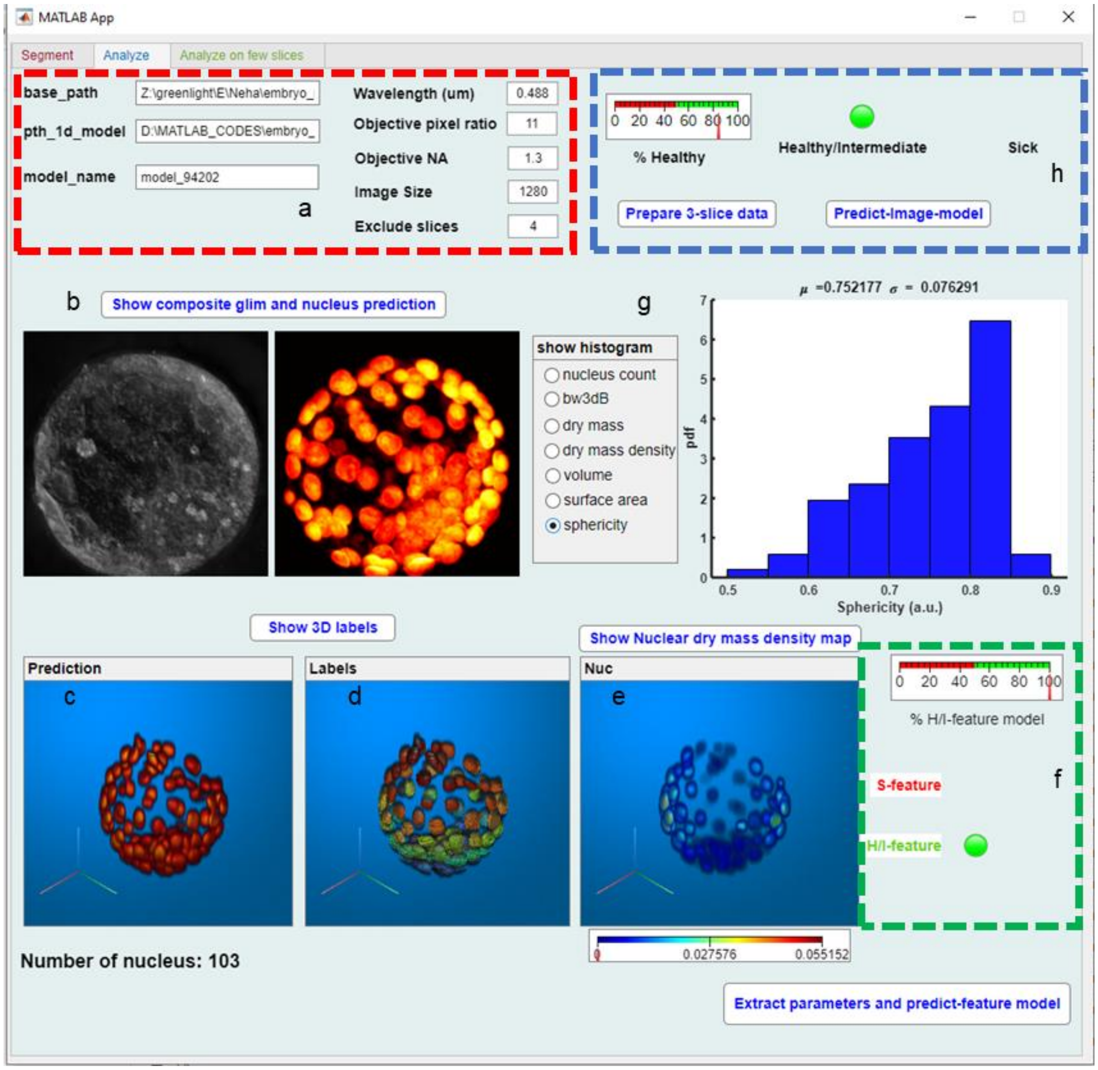


**Supplementary Figure S7. MATLAB standalone app panel 2: Analyze**. a. User input, b. Composite LS-GLIM and nucleus prediction c. 3D stacked nucleus prediction d. 3D segmented labels, e. Mean dry mass density map per nucleus, f. Health grading by FBM, g. Histograms of extracted features, h. Health grading by IBM.

**Analyze on few slices Panel**

This panel is a demo of sparse predictions feature of the IBM on live embryo data. Inputs in this panel (Fig. S8a) are:

1. **pth_to_base:** system path to the live embryo data
2. **prediction_filename:** IBM output csv file name
3. **ground_truth_filename:** name of the file containing z-slice names from which z position is extracted.
4. **glim folder:** folder containing 3 channel z-slice images to show the composite images (Fig. S8b)

Show composite button shows a montage of z-slice images of an embryo in steps of 3 µm. The z-slices positions are mentioned on the top of every tile (Fig. S8b). Random selections of z-slice positions can be made through the check box panel (Fig. S8c). When the required number of z-slice positions are selected (Fig. S8d), the Done switch should be turned 'On' to enable embryo-level prediction. The result is shown against the Status text and the percentage of z-slices in favor of the predicted class is displayed on the slider (Fig. S8e). To restart the process, the restart switch should be turned to On position.

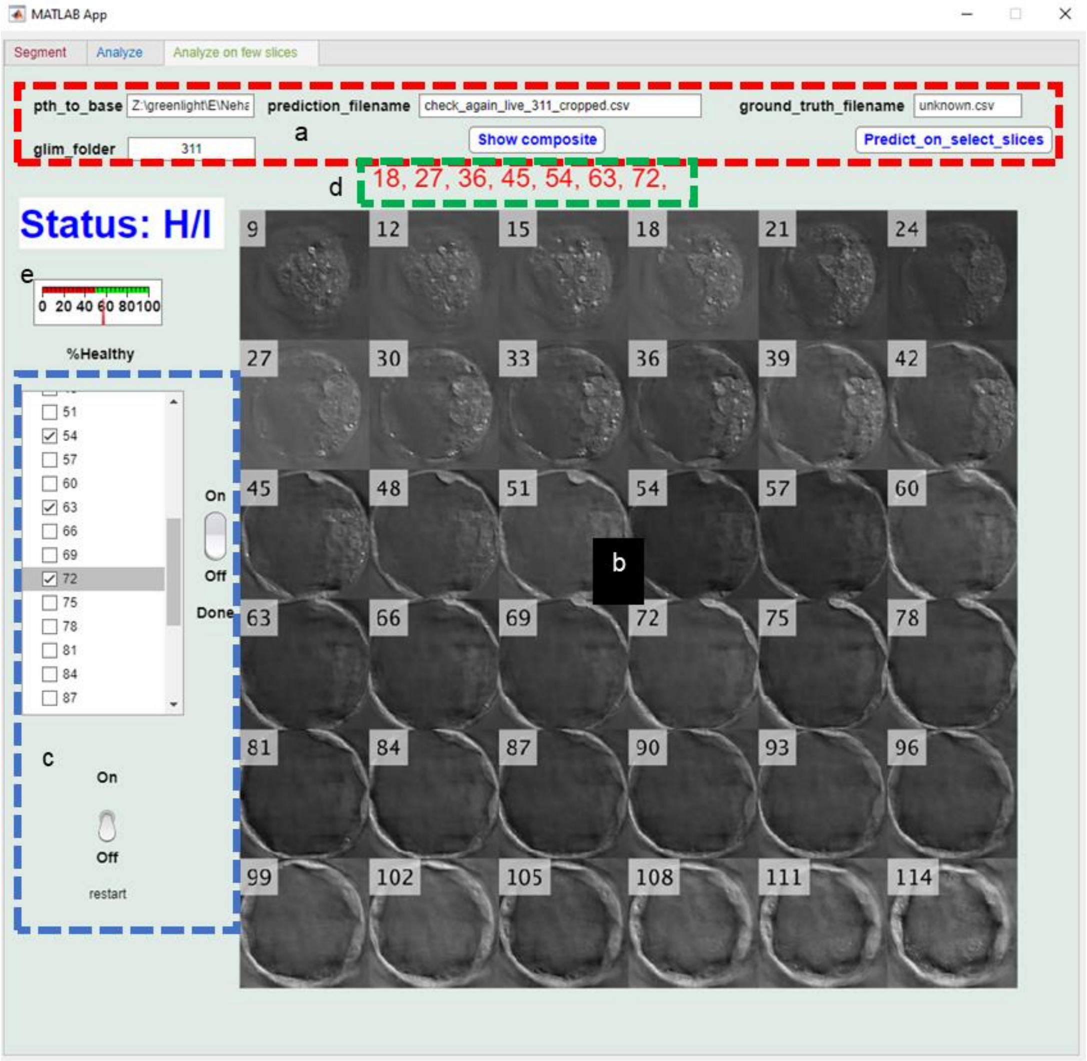


**Supplementary Figure S8. MATLAB standalone app panel 3: Analyze on fewer slices:** a. User input, b. Montage of z-slices of the individual embryo, c. Slice selection menu d. Selected z-slices, e. Health status by IBM on selected slices.

| name | actual_class | IBM_prediction | FBM_prediction |
|---|---|---|---|
| h4_d10_f2 | H/I | H/I | H/I |
| h4_d1_f2 | H/I | H/I | H/I |
| h4_d1_f4 | H/I | H/I | H/I |
| h4_d5_f2 | H/I | H/I | H/I |
| h4_d7_f2 | H/I | H/I | H/I |
| h4_d8_f3 | H/I | H/I | H/I |
| h4_d8_f4 | H/I | H/I | H/I |
| h4_d9_f1 | H/I | H/I | H/I |
| h5_d10_f3 | H/I | H/I | H/I |
| h5_d11_f4 | H/I | H/I | H/I |
| h5_d2_f1 | H/I | H/I | H/I |
| h5_d5_f1 | H/I | H/I | H/I |
| h5_d5_f2 | H/I | H/I | H/I |
| h5_d6_f1 | H/I | H/I | H/I |
| h5_d6_f2 | H/I | H/I | H/I |
| h5_d6_f3 | H/I | H/I | H/I |
| h5_d7_f1 | H/I | H/I | H/I |
| h5_d8_f1 | H/I | H/I | H/I |
| h1_d2_f1 | H/I | H/I | H/I |
| h1_d2_f3 | H/I | H/I | H/I |
| h1_d3_f5 | H/I | S | H/I |
| h2_d2_f1 | H/I | H/I | H/I |
| h2_d3_f1 | H/I | H/I | H/I |
| h2_d3_f10 | H/I | H/I | H/I |
| h2_d3_f4 | H/I | H/I | H/I |
| h2_d5_f2 | H/I | H/I | H/I |
| h2_d5_f5 | H/I | H/I | H/I |
| h2_d5_f7 | H/I | H/I | H/I |
| h2_d5_f9 | H/I | H/I | H/I |
| h3_d2_f2 | H/I | H/I | H/I |
| h3_d6_f1 | H/I | H/I | H/I |
| s4_d10_f8 | S | S | S |
| s4_d1_f5 | S | S | S |
| s4_d1_f7 | S | S | S |
| s4_d9_f3 | S | S | S |
| s5_d11_f1 | H/I | H/I | H/I |
| s5_d1_f1 | S | S | S |
| s5_d4_f5 | S | S | S |
| s5_d6_f7 | S | S | S |
| s5_d8_f2 | S | S | S |
| s5_d8_f5 | S | S | S |
| s5b_d10_f1 | S | S | S |
| s5b_d11_f1 | S | S | S |
| s5b_d12_f1 | S | S | S |
| s5b_d13_f1 | S | S | S |
| s5b_d14_f1 | S | S | S |
| s5b_d1_f1 | S | S | S |
| s5b_d2_f1 | S | S | S |
| s5b_d3_f1 | S | S | S |
| s5b_d4_f1 | S | S | S |
| s5b_d5_f1 | S | S | S |
| s5b_d6_f1 | S | S | S |
| s5b_d7_f1 | S | S | S |
| s5b_d8_f1 | S | S | S |
| s5b_d9_f1 | S | S | S |
| s1_d1_f5 | H/I | H/I | H/I |
| s1_d2_f2 | S | S | S |
| s1_d4_f1 | S | S | S |
| s1_d5_f2 | S | S | S |
| s2_d1_f2 | H/I | H/I | H/I |
| s2_d1_f4 | H/I | H/I | H/I |
| s2_d2_f1 | H/I | H/I | H/I |
| s2_d2_f4 | H/I | H/I | H/I |
| s2_d3_f8 | H/I | H/I | H/I |
| s2_d4_f4 | H/I | H/I | H/I |
| s3_d1_f3 | H/I | H/I | H/I |
| s3_d3_f1 | H/I | H/I | H/I |
| s3_d5_f2 | S | S | S |
| s3_d5_f4 | H/I | H/I | H/I |
| s3_d6_f4 | H/I | H/I | H/I |
| s3_d7_f1 | H/I | H/I | H/I |
| s3_d7_f3 | H/I | H/I | H/I |

**Supplementary Table 1**. Test performance on common in-distribution data of 72 fixed embryos.

| name | actual_class-exp1 | actual_class-exp2 | IBM | FBM |
|---|---|---|---|---|
| 211 | H/I | H/I | H/I | H/I |
| 212 | H/I | H/I | H/I | H/I |
| 213 | H/I | H/I | H/I | H/I |
| 221 | H/I | S | S | H/I |
| 222 | H/I | H/I | S | H/I |
| 223 | S | S | S | S |
| 311 | H/I | H/I | H/I | S |
| 312 | H/I | H/I | H/I | H/I |
| 313 | S | S | S | S |
| 314 | S | S | S | S |
| 321 | H/I | H/I | H/I | H/I |
| 322 | H/I | H/I | H/I | H/I |
| 323 | H/I | H/I | H/I | H/I |
| 324 | S | S | S | S |
| 111 | H/I | H/I | H/I | H/I |
| 121 | H/I | H/I | H/I | S |
| 122 | H/I | I/S | S | S |
| 123 | S | S | S | S |
| 124 | S | S | S | S |
| 131 | H/I | S | S | S |
| 141 | S | S | S | S |
| 161 | S | S | S | S |
| | | ACCURACY (Excluding expert decision mismatch) | 0.947 | 0.895 |

**Supplementary Table 2.** Test performance on out-of-distribution data of 19 time-instances of 8 embryos

| Feature | P-val (Normality) Lilliefors Test (alpha=0.05) | | | KW-statistics (alpha=1e-3) | P-val (Dunn-Multiple comparison) (alpha=1e-3) | | | Sample-size |
|---|---|---|---|---|---|---|---|---|
| | H | I | S | P-val | HI | IS | HS | [H,I,S] |
| **dm** | 0.0009 | 0.0009 | 0.0009 | 0.79e-54 | 0.45 | 0.11e-50 | 0.23e-43 | [2775,3827, 1186] |
| **dmd** | 0.0009 | 0.0009 | 0.0009 | 0.82e-73 | 0.23 | 0.18e-63 | 0.22e-64 | [2775,3827, 1186] |
| **volume** | 0.0009 | 0.0009 | 0.0009 | 0.002 | 0.012 | 0.012 | 0.544 | [2775,3827, 1186] |
| **surface_area** | 0.0009 | 0.0009 | 0.0009 | 0.19e-5 | 0.1 | 0.88e-6 | 0.38e-3 | [2775,3827, 1186] |
| **sphericity** | 0.0009 | 0.0009 | 0.0009 | 0.46e-19 | 0.55e-4 | 0.18e-10 | 0.12e-19 | [2775,3827, 1186] |
| **Nuc_count** | 0.69 | 0.17 | 0.77 | 0.36e-14 | 0.76 | 0.60e-12 | 0.27e-11 | [46,65,41] |
| **bw3dB** | 0.102 | 0.49 | 0.53 | 0.25e-15 | 0.82 | 0.51e-13 | 0.47e-12 | [46,65,41] |

| Feature | Levene test P-val | One-way ANOVA P-val | P-val (post-hoc Student T test) (alpha=1e-3) | | |
|---|---|---|---|---|---|
| | | | HI | IS | HS |
| **Nuc_count** | 0.399 | 0.14e-20 | 0.63 | 0.17e-17 | 0.25e-14 |
| **bw3dB** | 0.002 | 0.17e-21 | 0.82 | 0.63e-18 | 0.22e-17 |

**Supplementary Table 3.** Statistical test results

| Model Number | Model Type | Model Information | Status | Accuracy % (Test) | Accuracy % (Live) |
|---|---|---|---|---|---|
| 2.29 | Neural Network | Bilayered [10,10] | Tested | 98.17 | 60.39 |
| 2.7 | Naive Bayes | Gaussian Naïve Bayesian | Tested | 93.01 | 58.13 |
| 2.27 | Neural Network | Medium [25] | Tested | 98.12 | 56.73 |
| 2.15 | KNN | Fine KNN | Tested | 97.44 | 55.76 |
| 2.28 | Neural Network | Wide 100 | Tested | 97.86 | 54.68 |
| 2.4 | Discriminant | Linear | Tested | 97.86 | 54.68 |
| 2.26 | Neural Network | Narrow 10 | Tested | 98.54 | 54.14 |
| 2.3 | Neural Network | Trilayered [10,10,10] | Tested | 99.32 | 53.82 |
| 2.9 | SVM | Linear | Tested | 99.32 | 53.71 |
| 2.6 | Logistic Regression | Logistic | Tested | 98.54 | 53.07 |
| 2.11 | SVM | Cubic | Tested | 99.06 | 51.45 |
| 2.2 | KNN | Weighted | Tested | 98.23 | 50.91 |
| 2.5 | Discriminant | Quadratic | Tested | 96.56 | 50.91 |
| 2.8 | Naive Bayes | Kernel-Gaussian | Tested | 95.04 | 50.81 |
| 2.13 | SVM | Medium Gaussian | Tested | 99.69 | 50.38 |
| 2.12 | SVM | Fine Gaussian | Tested | 97.60 | 49.84 |
| 2.23 | Ensemble | Subspace Discriminant | Tested | 98.43 | 49.84 |
| 2.18 | KNN | Cosine | Tested | 98.33 | 49.84 |
| 2.14 | SVM | Coarse Gaussian | Tested | 99.63 | 48.98 |
| 2.3 | Tree | Coarse Tree | Tested | 98.17 | 48.87 |
| 2.16 | KNN | Medium | Tested | 98.80 | 48.87 |
| 2.17 | KNN | Coarse | Tested | 99.79 | 48.55 |
| 2.19 | KNN | Cubic | Tested | 98.75 | 48.33 |
| 2.32 | Kernel | Logistic regression kernel | Tested | 94.73 | 48.22 |
| 2.24 | Ensemble | Subspace KNN | Tested | 95.67 | 47.47 |
| 2.1 | Tree | Fine Tree | Tested | 97.03 | 46.18 |
| 2.2 | Tree | Medium Tree | Tested | 100.00 | 46.18 |
| 2.25 | Ensemble | RUSBoosted Tree | Tested | 98.17 | 46.18 |
| 2.21 | Ensemble | Adaboosted Tree | Tested | 100.00 | 46.18 |
| 2.1 | SVM | Quadratic | Tested | 99.84 | 44.67 |
| 2.31 | Kernel | SVM Kernel | Tested | 97.60 | 43.92 |
| 2.22 | Ensemble | Bagged Trees | Tested | 98.17 | 40.37 |

**Supplementary Table 4.** Initial model selection results-MATLAB
